# Structural analysis of the bacterial amyloid curli

**DOI:** 10.1101/2022.02.28.482343

**Authors:** Mike Sleutel, Brajabandhu Pradhan, Han Remaut

## Abstract

Two decades have passed since the initial proposition that amyloids are not only (toxic) byproducts of an unintended aggregation cascade, but that they can also be produced by an organism to serve a defined biological function. That revolutionary idea was borne out of the realization that a large fraction of the extracellular matrix that holds Gram-negative cells into a persistent biofilm is composed of protein fibers (curli; tafi) with cross-β architecture, nucleation-dependent polymerization kinetics and classic amyloid tinctorial properties. The list of proteins shown to form so-called ‘functional’ amyloid fibers *in vivo* has greatly expanded over the years, but detailed structural insights have not followed at a similar pace, in part due to the associated experimental barriers. Here we combine extensive AlphaFold2 modelling and cryo-electron transmission microscopy to propose an atomic model of curli protofibrils, and their higher modes of organization. We uncover an unexpected structural diversity of curli building blocks and fibril architectures. Our results allow for a rationalization of the extreme physico-chemical robustness of curli, as well as earlier observations of inter-species curli promiscuity, and should facilitate further engineering efforts to expand the repertoire of curli-based functional materials.

## Introduction

Although once considered an implausible target for structural biology, recent developments in cryo-electron microscopy (cryoEM) and helical processing have facilitated the structure determination of amyloid fibrils at near atomic resolution ^1^. A plethora of experimentally determined structures of both *in vitro* generated and *ex vivo* isolated amyloids has revealed a bewildering structural diversity of fiber architectures, including the concept of fiber polymorphs or “strains” of otherwise identical or closely related sequences ^2-5^. Despite differences in proto-fibril architecture and fiber helical symmetry, most amyloid structures share several conserved features that allow us to expand the definition of the amyloid fold beyond the traditional “cross-beta” adage. To the best of our knowledge, all currently described amyloid structures consist of a repetitive stacking of intricate, serpentine, planar β-strand arrangements that are stabilized by steric zipper motifs wherein interdigitated residue side chains make extensive Van der Waals, electrostatic, hydrophobic and hydrogen bonding contacts. Axial stacking of planar peptides by strand-strand docking and β-arcade formation drive protofibril formation, usually followed by the helical winding of multiple protofibrils into a remarkably stable helical superstructure. It is precisely this helical symmetry that has been leveraged with great success in modern cryoEM approaches to resolve the structural details of a wide range of amyloid structures.

Most of those structures belong to a subfamily of amyloid species that are correlated to a host of (neuro)degenerative, systemic deposition and misfolding diseases. For that reason, those amyloidogenic proteins are colloquially referred to as pathological amyloids (PAs). These disease-associated amyloids have in common that they represent a non-functional and off-pathway misfolding and aggregation event of proteins or protein fragments destabilized from reaching their native structure by mutation, environmental conditions or misprocessing. There is a second branch of the amyloid family, found across all domains of life, which consists of proteins that evolved to fulfill dedicated biological roles (such as adhesion, storage, scaffolding, etc.) by adopting the amyloid state – affording these proteins the term functional amyloids (FAs) ^6-8^. Like pathological amyloids, FAs show nucleation-dependent aggregation into fibers with cross-β characteristics. An enigmatic question is whether FA pathways include selected traits to mitigate or lack the cytotoxic gain of function properties so commonly associated with pathological amyloid depositions. For bacterial amyloid pathways like curli and Fap, it is clear that accessory proteins ensure a timely and localized amyloid deposition, including chaperone-like safeguards that prevent or stop premature amyloidogenesis ^9,10^. Whether the structures of the FA subunits and fibers also include adaptive traits that lower cytotoxicity is much less understood, however. Interestingly, there are indications from *in vitro* experiments that FAs produced by various pathogenic bacteria can exhibit cross-reactivity with PAs ^11^, and reports of a potentially infectious induction or aggravation of pathological amyloid depositions by direct cross-seeding or indirect (inflammatory) effects in humans and animals exposed to bacterial amyloids ^12,13^. As there are only a few structures available for FAs ^14,15^, it is unclear at this point to what extend FAs are structurally related to PAs, and if they form a separate branch of amyloid architectures. Nor is it clear what the molecular mechanism could be for this inter-species FA/PA amyloid promiscuity.

The amyloid fold is reported as one of the most stable quaternary protein states. The pre-programmed self-assembly properties of FAs have attracted general interest for their use in synthetic biology and biotechnology applications precisely because of their extreme robustness and ability to spontaneously develop with minimal intervention or catalytic assistance. Although some successes have been reported in the development of advanced functional materials derived from FAs ^16,17^, it stands to reason that the field will benefit from a more profound structural understanding to inform a set of rational design principles. Given the recent advances in *de novo* protein design we believe this could help usher in an era of tailor-made amyloid polymer production.

To address the structural biological blind spot with regards to FAs, we pursued the structure of the model bacterial FA curli. Curli fibers are a major component of the extracellular matrix of Gram-negative bacteria and are expressed under biofilm forming conditions where they fulfill a scaffolding and cementing role in the extracellular milieu to reinforce the bacterial community ^18,19^. The major subunit of *Escherichia coli* curli is the 13.1kDa pseudo-repeat protein CsgA, which is secreted as a disordered monomer that forms cross-β fibers with classic amyloid tinctorial properties under a wide range of conditions ^20^. In its native context, formation of cell-associated CsgA fibrils requires the minor curli subunit CsgB, which in turn is bound to the outer membrane CsgG-CsgF secretion pore complex ^21,22^. Although structural models have been proposed for a folded CsgA monomer based on homology modelling ^23^, solid-state NMR ^24^ and co-evolutionary coupling analysis ^25^, there is no available experimental curli structure. Here we present an in-depth bio-informatics study of the amyloid curli core by analyzing the primary sequences of a local CsgA catalogue that was constructed through exhaustive mining of the bacterial refseq database. Based on that analysis we propose different structural classes of curli subunits, and discuss the expected implications for the CsgA fold, and by extension the curli fiber architecture. We test and validate those predictions based on extensive AlphaFold 2 (AF2) modelling of 2500+ CsgA homologue sequences. Next, we selected two CsgA candidates for detailed cryoEM analysis, from which we derive distance restraints that are in excellent agreement with the proposed AF2 models. Based on this, we present a molecular model for a CsgA protofibril and discuss its modes of higher-order organization in the extracellular matrix and fibers formed *in vitro*.

## Results

### Variations in the number, consistency, and length of curli repeats across the Gram-negative bacteriome

First, we setup a local database of homologue CsgA sequences by mining the Refseq bacterial genome repository using the HMM curli profiles constructed by Dueholm *et al* ^19^. For CsgA in particular, we use an HMM profile that is specific to the amyloid repeat, and therefore does not differentiate between the major curlin subunit CsgA, and the minor curlin subunit and purported nucleator CsgB. In what follows, we will refer to this group of curli repeat-containing proteins as CsgA, bearing in mind that a subpopulation will correspond to CsgB sequences.

From a total of 201210 bacterial genomes that were searched, 43279 (22%) genomes contained one or more predicted CsgA sequences that harbor one or more curli repeat signatures (Fig.1a). This correlated well with the presence of the other genes in the curli operon. From the 43279 genomes that contained CsgA, we detected CsgG, CsgF, CsgE and CsgC or CsgH in 41041, 41597, 41090 and 37945 or 1839 genomes, respectively. Only a small minority (1033) of the CsgA containing genomes lacked a copy for curli secretion channel CsgG. This low number of CsgA orphans, combined with the excellent correlation to the presence of the other curli biogenesis proteins, demonstrates that the resulting dataset of CsgA protein sequences is predominantly derived from *csga* genes that are embedded in a curli operon. The vast majority (40559) of these genomes contains two CsgA-like copies, which likely to correspond to the functional diversification into CsgA and CsgB^26^. Interestingly, genomes can be found with more than two CsgA homologues, with the number of CsgA copies per genome following an exponential decline towards 14 copies and a maximum detected of 30 (GCF_003076275.1_ASM307627v1) (Fig.1a). After removal of partial entries, we obtain a list of 87205 putative CsgA sequences, which reduced to a dataset of 8079 unique mature domain CsgA sequences upon removal of duplicates and truncation of the leader sequences. As an initial proxy for the number of curlin repeats per CsgA homologue, we use the reported number of hits for the curlin repeat in the HMMER log-file. This yields a distribution of CsgA homologues with repeat numbers that span the spectrum from 4 to 62 (WP_189563214.1), with local maxima at 5 or 7-8 curlin repeats (the former typical of Enterobacteria like *Escherichia* and *Salmonella*), and secondary maxima around 15 and 22 (e.g. Pseudomonas; Fig.1b). By plotting the number of predicted repeats versus the length of the primary sequence, the average length of a curlin repeat is found to be 23±5 (Fig.1b inset). The low standard deviation demonstrates that the average length of a single curlin repeat is strongly conserved for CsgA, and consequently so will be the lateral dimensions of the resulting fibers (see below).

**Fig. 1:**
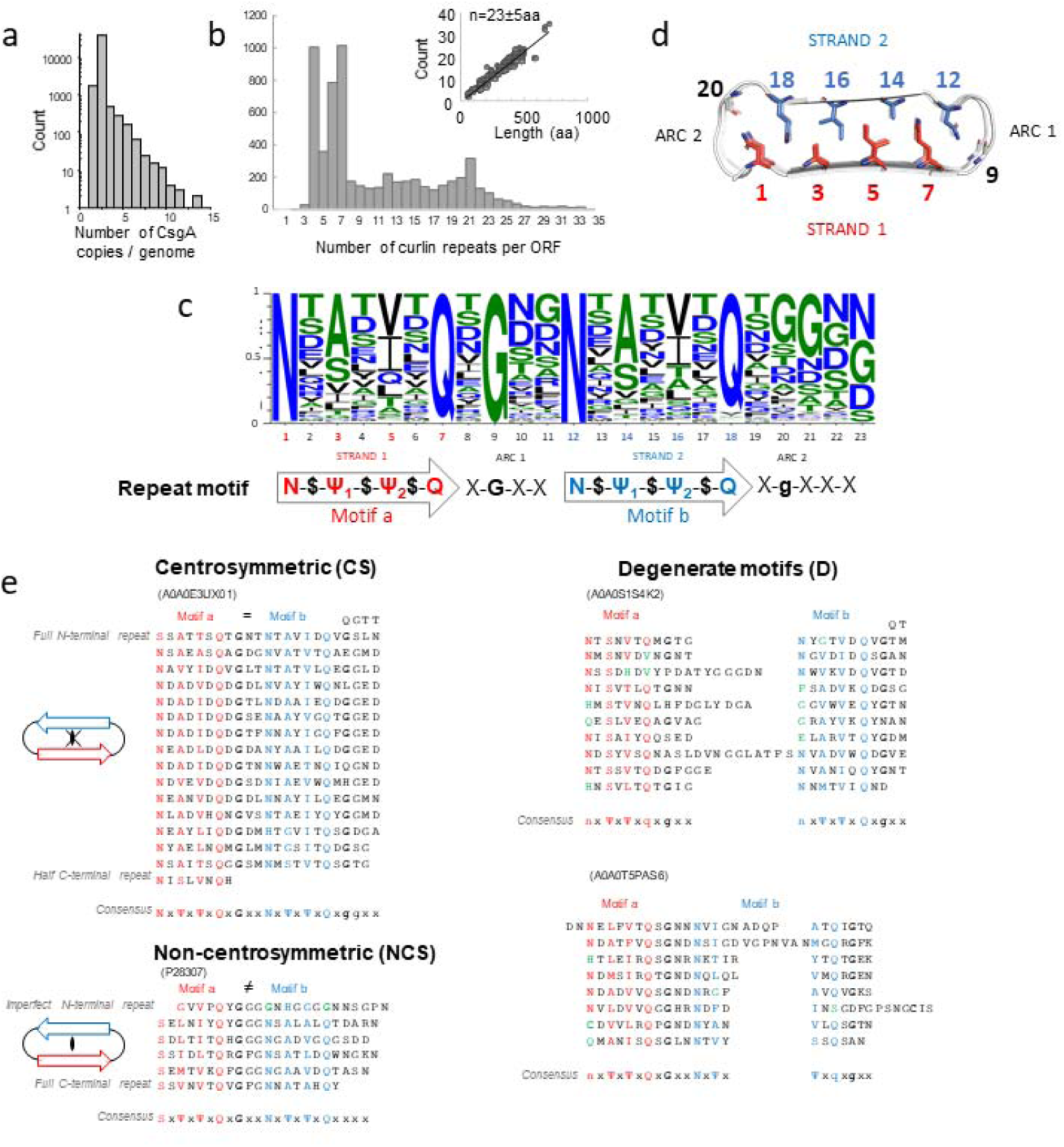
Primary sequence analysis of CsgA homologues. **(a)** Distribution of the number of csga-homologues detected per genome; **(b)** Distribution of the number of predicted curlin repeats. Inset: number of curlin repeats per protein length, with a slope of n=23±5 aa; **(c)** Consensus sequence of a curlin repeat with the proposed curlin motif denoted underneath; **(d)** On-axis view of a β-solenoid fold with conserved inwards facing residues highlighted; **(e)** Representative primary sequences (Seq ID in parentheses) of three sequence classes of CsgA, stacked according to their predicted curlin repeats, with respective consensus motifs indicated below (colored red and blue for motif A and B, resp., green when deviating).

The large variation in CsgA repeat numbers prevents investigation of sequence conservation and variability in the amyloid core via global sequence alignment. Rather, we extracted 53574 non-overlapping, linear segments of 24aa that are centered on a QX_10_Q motif (and permutations thereof) which represents the prototypical amyloid kernel of the CsgA curlin repeats. From this we derive a consensus sequence ^27^ that exhibits remarkable conservation at key sites and shows curlin repeats can be divided into closely related motifs A and B, typically separated by a 4-residue sequence with X-G-X-X signature, where X is any amino acid (Fig. 1c). In the predicted CsgA structure (see below) these curlin repeat elements form strand 1, 2 and the connecting loop or ‘β-arc’ of a β-arch as defined by Henetin et al. ^28^. As such, the residues indicated in red and blue below the WebLogo and the structural representation (Fig.1c, d) map, respectively, to the inward facing steric zipper residues and outward facing surface residues of the strand-arc-strand (i.e. β-arch) motif that constitutes a single curlin repeat. The base signature of the motifs A and B consists of N-$-Ψ_1_-$-Ψ_2_-$-Q where Ψ_n_ are hydrophobic residues (A,I,V,L,F), S or T and point to the inside of the β-arch, together with a well conserved N at the start and Q and the end of the motifs. Residues on locations corresponding to $ are surface exposed and are predominantly polar or charged (T,S,D,E,Q,N,R,Y), suggesting that most curli fibers can be expected to be hydrophilic. Motif B is followed by 4-5 residues, most typically in a X-G-X-X-(X) motif. This loop region forms a second β-arc, connecting motif B to motif A in the subsequent curlin repeat (Fig. 1d).

When analyzing the curli subunits in further detail, we identify three sequence classes: centrosymmetric (CS), non-centrosymmetric (NCS), and degenerate (D) (Fig. 1e). The CS-class contains curlin repeat sequences for which the core-facing residues in motif A (i.e. N, Ψ_1_, Ψ_2_ and Q at positions 1, 3, 5 and 7, resp.) form a (nearly) identical repeat in motif B (i.e. positions 12, 14, 16 and 18), with A0A0E3UX01 from *Pontibacter korlensis* as representative example (Fig.1e). Considering the expected tertiary structure (Fig.1d), this yields a steric zipper that is centrosymmetric in the placement of the core-facing residues. Notably, outward facing residues (i.e. $) do not adhere to the selective pressure that retains centrosymmetry seen in the curlin repeats of CS-class CsgA sequences. NCS-class CsgA sequences are composed of curlin repeats for which the core-facing residues in motifs A and B are not (near) perfect repeats, which will develop steric zippers that are non-centrosymmetric, as is the case for CsgA from *E. coli* or *Salmonella enterica* (Fig.1e). In EcCsgA, N is substituted by S in motif A, and positions Ψ_1_, Ψ_2_ are bulky hydrophobics (I, L, M), versus mostly small hydrophobics (A, V) in motif B. A third class of CsgA sequences can be discerned, showing degenerate repeats, i.e. repeats for which either motif A and/or B is incomplete (e.g. absence of the flanking N in motif A or Q in motif B), and/or where curlin repeats are considerably longer than the average 23 residues as a result of insertions downstream of motif A (i.e. arc 1) or motif B (i.e. arc 2), with A0A0S1S4K2 of *Sediminicola sp. YIK13* as a representative example. Less commonly, insertions can also fall inside motif A or B, however, with A0A0T5PAS6 of *Roseovarius indicus* as a representative example (Fig. 1e).

Finally, our sequence analysis reveals that CsgA sequences can terminate in a full or half curlin repeat, i.e. terminate in a prototypical motif A – arc – motif B sequence or in a motif A sequence only (Fig. 1e). Since motif A and B represent strand 1 and 2 of the curlin β-arches, the termination in a complete or incomplete β-arch can be expected to influence intersubunit contacts in the resulting fibers (see below). Of further notice is that CsgA sequences can start with a single degenerate curlin repeat where the flanking N(S) and Q of both motif A and B are mutated, frequently to G, with EcCsgA (Fig. 1e) as representative example. In case of EcCsgA, this degenerate N-terminal repeat is known as N22 and represents a targeting sequence that binds the CsgG secretion channel and that was found not to incorporate into the amyloid core of the curli fiber ^20,29^. Of note, our sequence analysis indicates the presence of an N22-like sequence is an outlier more than the rule. In the following section, we first discuss the predicted tertiary structures for CS-, NCS- and D-class CsgA sequences and the structural implications for the folded CsgA monomer. Note that we focus here on a structural classification of curlin motifs and subunits, for a phylogenetic analysis we refer the readers to earlier work by Dueholm *et al* ^19^.

### AlphaFold2 modelling of 2500+ CsgA homologues reveals a canonical β-solenoid and a conserved amyloid kernel

We employed the *localcolabfold* implementation ^30-32^ of AlphaFold2 (AF2) to predict the structure of 2686 unique CsgA sequences. This constitutes a representative subset of the total CsgA database, encompassing the full range of predicted repeats as well as the diversity in repeat motifs. We obtain an average pLDDT value of 83±9 for the total dataset (see Supporting Fig. 1). For 741 models (27%), the pLDDT value is higher than 90, whereas 237 models (8%) score below 70. DeepMind reports pLDDT > 90 as high accuracy predictions, between 70 and 90 as good backbone predictions, and pLDDT < 70 as low confidence and to be treated with caution ^31^. All models share a similar β-solenoid architecture, wherein curlin repeats fold into strand-β-arc-strand motifs that stack vertically to produce an in-register double, parallel β-sheet structure with a single strand stagger (i.e. 2.4 Å) between both sheets (Fig. 2, Supporting Fig.2). This topology fits with our cryoEM observations on mature curlin fibrils, which we discuss in detail below. It also means that variations in the number of repeats correlate linearly to the long-axis dimension of a CsgA monomer, and that the curliome spans a continuum of solenoidal monomers.

**Fig. 2:**
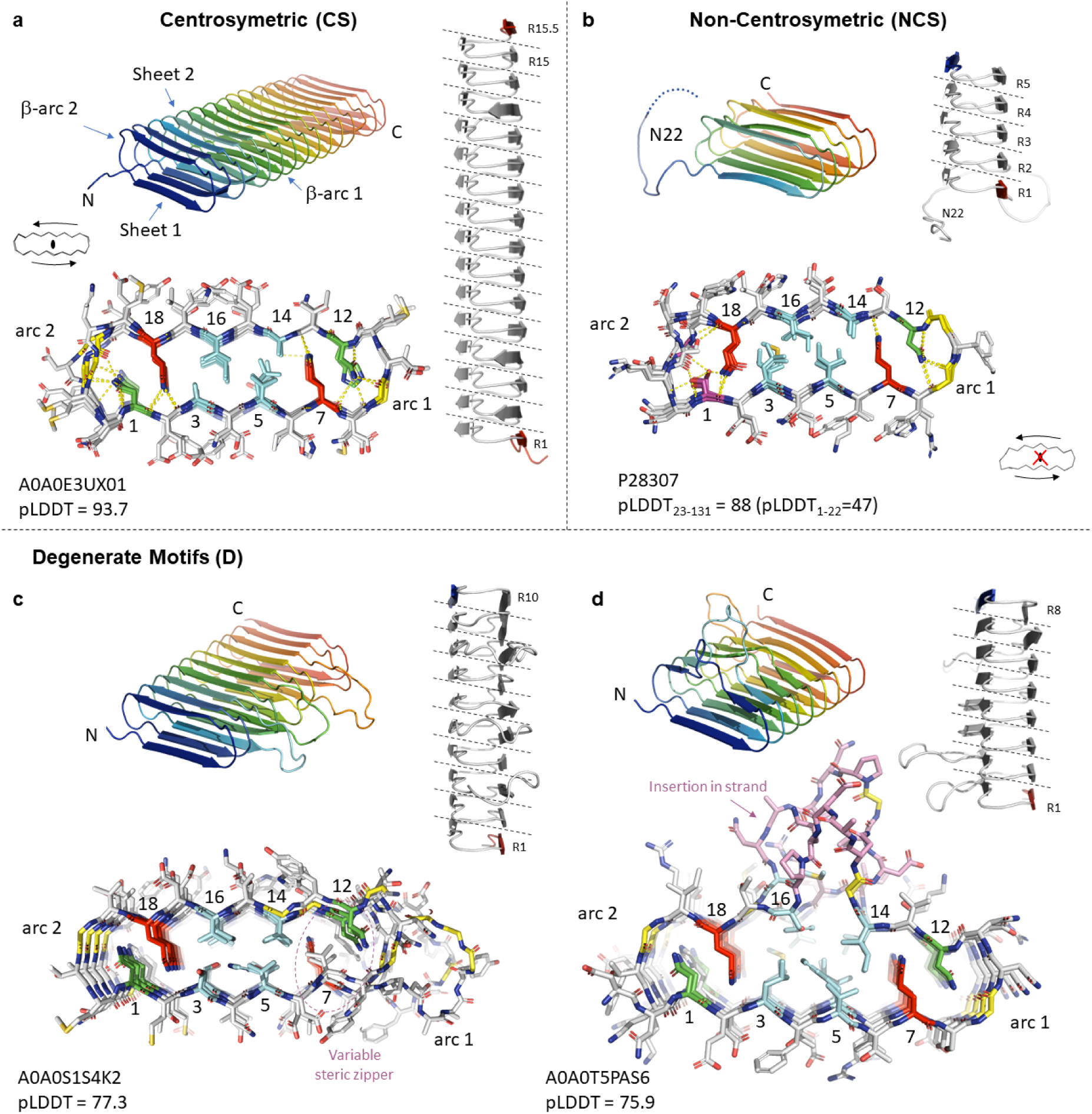
Structural classification of curlin subunits as predicted by AF2. Cartoon representation of representative examples (with Uniprot accession ID and AF2 pLDDT score) for each category, with a transversal view shown below in stick representation to highlight the steric zipper residues and the alignment of the repeats along the long axis. All models share the same basic architecture, i.e helical assembly of repeating strand-turn motifs, resulting in a β-pleated core flanked by two β-arcades. **(a)** AF2 model of R15.5 from *Pontibacter korlensis*, characterized by a centro-symmetric zipper motif and both N- and C-terminal strands on the same sheet of the β-solenoid; **(b)** CsgA from *E*.*coli* with terminating strands on opposing sides of the solenoid, and a non-centrosymmetric motif as well as a disordered N-terminus (1-22); **(c)** CsgA from *Sediminicola sp. YIK13* with variable steric zipper residues leading to degenerate core motifs and arc2 insertions; **(d)** CsgA from *Roseovarius indicus* with loop and strand insertions of variable length.

As a representative structure of CS-class CsgA monomers we discuss the AF2 model of A0A0E3UX01 (pLDDT=93.68; Fig.2a), which has 15 curlin repeats and an additional half repeat at the C-terminus (hereafter referred to as R15.5). AF2 predicts a highly regular (the average RMSD between consecutive repeats is 0.18±0.02 Å), left-handed β-solenoid with a negligible twist. Thus, the monomer consists of an almost pure translational stacking of the β-arches formed by the curlin repeats. A transversal, on-axis view provides further insight into the structural nature of the steric zipper, which comprises a central hydrophobic core composed of the Ψ_1_ and Ψ_2_ residues in motif A and B, flanked on either side by a glutamine (Gln), followed by an asparagine (Asn), forming the centrosymmetric core predicted based on motif A and B sequence features. The side chain amide groups of Gln 7 and 18 in the curlin core (the numbering as in consensus motif Fig. 1c) engage in extensive hydrogen-bonding network, forming a ladder of H-bonds with the equivalent Gln in the flanking repeats, and with the main-chain carbonyls of the opposing strands belonging to either its parent or adjacent repeat at positions 13 or 2, respectively (Supplementary Fig. 3a). These interactions are facilitated by the stagger between the two sheets and likely form a major contribution to the stabilization of the sheet-on-sheet packing. Ψ_1_ and Ψ_2_ residues in motif A and B interdigitate into a hydrophobic core, likely providing additional stabilization of the sheet packing in the β-solenoid. Of note, our HMM search identified CS-class CsgA-like sequences where Ψ_1_ and Ψ_2_ form a purely hydrophilic core consisting of Ser / Thr, such as A0A249PTX6 of *Sinorhizobium fredii* (Supplementary Fig. 3b). Although these sequences consist of regular repeats of the canonical N-$-Ψ_1_-$-Ψ_2_-$-Q-X-G-X-X-N-$-Ψ_1_-$-Ψ_2_-$-Q-X-G-X-X kernel and are predicted to adopt the curli β-solenoid, these did not form part of a *Csg* operon.

Whereas the Glns stabilize inter-repeat interactions, as well as cross-strand packing, N1 and N12 residues form an extensive H-bond network stabilizing β-arc 2 and β-arc 1, respectively. Asn 1 forms 4 mainchain H-bond interactions with residues G20, X21, X23 in the +2 curlin repeat and N1 of the +1 repeat, whilst N12 H-bonds the mainchain of X8, G9, X10 and N12 in the +1 repeat. Despite low sequence conservation in the positions 10-11 across the 15 repeats, AF2 predicts the main chain trace to be quasi isomorphous throughout (average rmsd of 0.04±0.01 Å for equivalent C-α positions in the β-arches), yielding an uninterrupted β-arcade that is further stabilized by intra-main chain H-bonds in consecutive β-arches. If we normalize all detected polar contacts to the number of repeats, then R15.5 forms 29.6 H-bonds per repeat.

As a representative structure of NCS-class CsgA monomers, the AF2 prediction for CsgA from *E. coli* (P28307; pLDDT_23-131_=88; Fig.2b) is a left-handed, 5 repeat, β-solenoid with 25.4 H-bonds per repeat. As expected from the sequence analysis, packing of the motif A and motif B β-sheets (sheet 1 and 2) loses the centrosymmetry seen in CS-class CsgA. In EcCsgA, the Asn column in position 1 is replaced by a Ser column, breaking the centro-symmetric nature of the steric zipper. This column of serine residues can be regarded as a functional replacement of N1 in that they also stabilize the β-arc 2, but have lower H-bonding potential with respect to asparagine. EcCsgA also loses centrosymmetry in the hydrophobic core, consisting of bulky hydrophobic Ψ_1_ and Ψ_2_ residues in motif A (position 3 and 5) versus small hydrophobics in motif B (14 and 16) (Fig 2b). This compares to a symmetric packing of the Ψ_1_ and Ψ_2_ positions of CS-class sequences (Fig. 2a). An additional loss of symmetry is manifested at the level of the β-arcs: arc 1 consists of the prototypical X-G-X-X motif (in EcCsgA X-G-X-G) and has a narrow curvature, whereas arc 2 is 4-5aa wide and shows poor sequence conservation. The average RMSD between consecutive repeats is 0.19±0.01 Å which is similar to the value obtained for R15.5. The N-terminus of EcCsgA holds an imperfect curlin repeat known to serve as secretion signal and shown to remain protease accessible in the mature fiber ^20,29^. The average pLDDT value for these first 22 residues (N_22_) is 47, and N_22_ is modelled inconsistently between various AF2 predictions, either disordered, or partially docked onto the β-solenoid scaffold. Finally, the EcCsgA solenoid terminates on a full-repeat which means the subunit terminates with a sheet 1 (motif A) and sheet 2 (motif B) overhang at N- and C-terminus, resp. In R15.5, which terminates in a half-repeat, both the N- and C-terminal overhang are located on sheet 1 (Fig. 2a).

As an example of a curlin monomer with degenerate repeats we focus on A0A0S1S4K2 for which AF2 predicts a left-handed, 10-stranded β-solenoid with an average 25.6 H-bonds per repeat (pLDDT=77.3; Fig.2c). The A0A0S1S4K2 sequence shows deviations in the N and Q columns of motif A and/or B, including substitutions for hydrophobic residues. These N/Q substitutions disrupt the regular H-bond enforced steric zipper, and are frequently found to flank positions of protrusions in the arc connecting the respective motifs. This translates in a moderate increase of the RMSD across the neighboring repeats (0.35±0.07 Å). A further departure from a symmetric β-solenoid fold continues for our fourth example. The AF2 model for A0A0T5PAS6 comprises a left-handed solenoid with 8 repeats, with insertions in the strand 1 of repeats 1, 2, 3, and 4 (pLDDT=75.9; Fig.2d). The net inter-repeat RMSD for equivalent Cα positions (i.e. disregarding these insertions) is 0.45±0.05 Å and reflects the local deviations from the ideal solenoid geometry to accommodate said insertions. This does not, however, translate in a reduction of the total H-bonding potential, which is 29.5 per repeat.

### Handedness of the curlin fold

Close inspection of the library of CsgA AF2 models revealed an ambiguity in handedness of the predicted β-solenoids. This ambiguity correlated with the dept of the multiple sequence alignments. When run without MSA or in case of sparsely populated MSAs (i.e. when using the default mmseq2 in colabfold) CsgA sequences were frequently predicted as right-handed β-solenoids, whereas the same sequences run through AF2 with well populated MSAs obtained using jackhmmer consistently lead to a left-handed β-solenoid. This is illustrated in Supporting Fig.4 for R15.5. Interestingly, both predictions have similar predicted alignment errors, contact maps and overall pLDDT (Right: 92.8; Left: 93.7) and pTMscores (Right: 0.91; Left: 0.89) precluding an *a piori* differentiation between both possibilities. MolProbity analysis ^33^ of the top-ranking, AMBER relaxed models for the L- and R-variant produced similar structural statistics overall, with the MolProbity score essentially indistinguishable (Right: 0.68; Left: 0.65). The most notable difference between both models -apart from the handedness-was the overall twist of the two sheets in the β-solenoid. Right-handed models (L) consistently have a ±20° twist, whereas right-handed (R) models do not (Supporting Fig.4).

In absence of kinetic selective factors of the respective folding pathways, it seems that the L- and R-model are both plausible theoretical end-states. Are curli fibers racemic mixtures in practice or is there a discriminating factor? Given the non-zero twist for the R-model, we predict that an R15.5 R-protofibril would be helical in nature, with a rise and twist of 7.2nm and 20° (i.e. 4.8 Å and 1.3° per repeat), respectively, whereas an L-protofibril would be constructed from translational symmetry only (see the following section for additional information as well as Supporting Fig.4). The former is in direct conflict with our cryoEM data (see below) as a helical R-protofibril would produce 2D class averages that span a continuum of fibril orientations, which is not observed experimentally, indicating R-protofibrils are very rare or absent from the dataset. Similar to R15.5, AF2(mmseqs2) produces a right-handed model for EcCsgA with an 11° twist (i.e. ∼2.2° per repeat), whereas the AF2(jackhammer) left-handed model shows a pure translation of the curlin repeat. The latter is supported by our cryoEM data on curlin fibers that were purified from a bacterial biofilm (discussed in detail below). Thus, at least for *P. korlensis* and *E. coli* CsgA, both *in vitro* and *ex vivo* curli fibers were found to (predominantly) consist of subunits comprising a left-handed β-solenoid.

### A conserved repeat architecture facilitates docking of disparate monomers into fibrils

Having established the structural hallmarks of a curlin monomer, we turn our attention to predictions of homo-and heteromeric assemblies. Curlin monomers end in two open-edged β-sheets with a 2.4 Å stagger, i.e. resulting in a single strand overhang at either end. Thus, subunits show a motif A (sheet 1) overhang at the N-terminus, and a motif B (sheet 2) or motif A (sheet 1) overhang at the C-terminus for sequences ending in a full or half curlin repeat, respectively (Fig. 3a). If the packing interface between two monomers can be predicted with reasonable confidence, then a model for the curli architecture can be inferred. To do this, we first benchmark the capabilities of AF2 to accurately predict and reproduce specific features of fibrous protein assemblies, such as the presence of a screw axis and fiber polarity. For this, we performed AF2 predictions of the bactofilin filaments of *Thermus thermophilus* which are composed of end-to-end associated β-helical domains. We use the filament cryoEM structure 6RIB as a reference and compare it to a trimer model (pLDDT=86.2; pTMscore=0.75) that was predicted using AF2 multimer (Supporting Figure 5). We note that the cryoEM structure was deposited on 2019-04-23, whereas AF2 is trained on protein chains in the PDB released before 2018-04-30. Overall, there is excellent agreement between the predicted and experimental structure, with an RMSD of 1.53Å. AF2 accurately predicts the apolar nature of the filaments (i.e. succession of head-to-head and tail-to-tail interfaces), the handedness of the β-solenoid, the disordered N- and C-terminal tail as well as the helical nature (i.e. presence of a screw axis). This reinforced our confidence in the proposed AF2 methodology for curlin modelling and prompted us to look at an AF2 CsgA trimer in further detail.

**Fig. 3:**
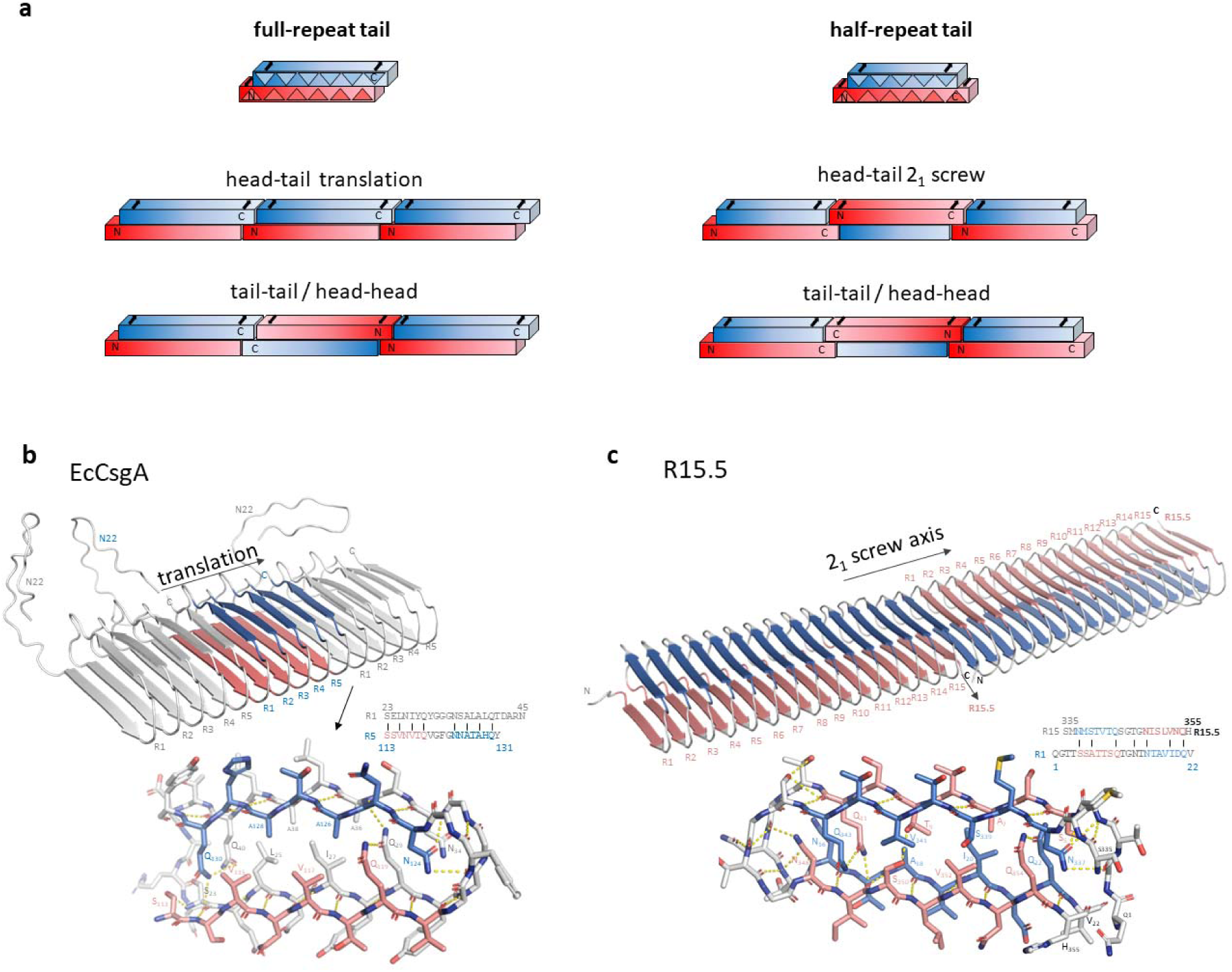
Predicting protofibril architecture using AF2. **(a)** Schematic representation of monomeric building blocks for CsgA sequences ending in a full or half repeat, creating a C-terminal motif B or motif A overhang, respectively. Head to tail or tail – tail / head-head stacking of CsgA monomers will result in a fiber with a cross-monomer stacking of the β-arcades. Head to tail stacking results in parallel β-arcade throughout the fiber, head-head / tail-tail alternates direction of the β-strands, creating an antiparallel interface. A head to tail stacking of CsgA sequences with half-repeat tail implies a 2_1_ screw axis for productive H-bonding of the β-arcade at the subunit interfaces. **(b, c)** Multimeric CsgA assemblies represent minimalistic protofibrils: **(b)** *Ec*CsgA trimer as predicted by AF2 wherein monomers stack ‘head-to-tail’ via a unique R5/R1 interface resulting in a polar protofibril (only translational symmetry) with an R1 and R5 terminus. Stick representation of the R5/R1 interface with putative inter-chain hydrogen bonds shown in dashed lines; For the central CsgA protomer, motif A and motif B are coloured red and blue, resp. **(c)** R15.5 dimer as predicted by AF2 wherein two monomers stack ‘head-to-tail’ and making a 180° rotation with respect to each other. The protofibril has one unique type of inter-molecular interface (R15.5/R1), with a 2_1_ translational symmetry and polar termini formed by repeats R1 and R15.5, respectively.

The AF2 model of an EcCsgA trimer consists of three head-to-tail stacked monomers that interact via their R5/R1 repeats by means of β-sheet augmentation, forming an extended solenoid fold (Fig.3b, Supporting Fig.5). This proto-fibril assembly is constructed from pure translational symmetry, with no screw-axis present or any measurable twist. Looking at the interface between two monomers, we find a near seamless transition of the β-solenoid hydrogen bonding network, i.e. the R5/R1 interface contains 23.3 putative H-bonds (averaged over the trimer) which is on par with the average number of H-bonds between the repeats within a single CsgA monomer. This continuity is facilitated by the low RMSD values between R1 and R5, as well as the conservation of the steric zipper residues leading to an inter-molecular digitation that is very similar to the intra-molecular contacts. The motifA-arc1-motifB-arc2 β-arcade continues uninterrupted. AF2 predicts N_22_ as disordered and excluded from the curli β-arcade, and thus protruding from the fibers, consistent with its reported proteolytic susceptibility in mature curli fibers ^20^.

It is worth pointing out that, although not predicted by AF2, CsgA might also be able to oligomerize by consecutive tail-to-tail / head-to-head interactions (Fig. 3a, Supporting Figure 6). To that end, we manually brought two CsgA molecules in R5/R5 contact, and performed a local docking using RosettaDock to optimize the local geometry (Interface score: -9.5). We reference this tail-to-tail dimer to a similarly optimized head-to-tail CsgA dimer where we used the AF2 model as the input to RosettaDock (Interface score: -15.5). Both models have a similar number of putative H-bonds at the dimer interface (23 *versus* 21), but the tail-to-tail model packs anti-parallel, triggering a lateral offset of 1 residue between both strands at the interface, and poor contacts in the N/Q steric zippers at the arcades as a result. If we extrapolate such a dimer to a protofibril, it is expected to produce a staggered pattern with a frequency of 2nm, which we do not observe experimentally (see Fig.4c), reaffirming the head-to-tail model produced by AF2. Furthermore, we previously found EcCsgA fibers to be polar, an observation that is only compatible with a head-to-tail interaction of the curli subunits ^34^.

**Fig. 4:**
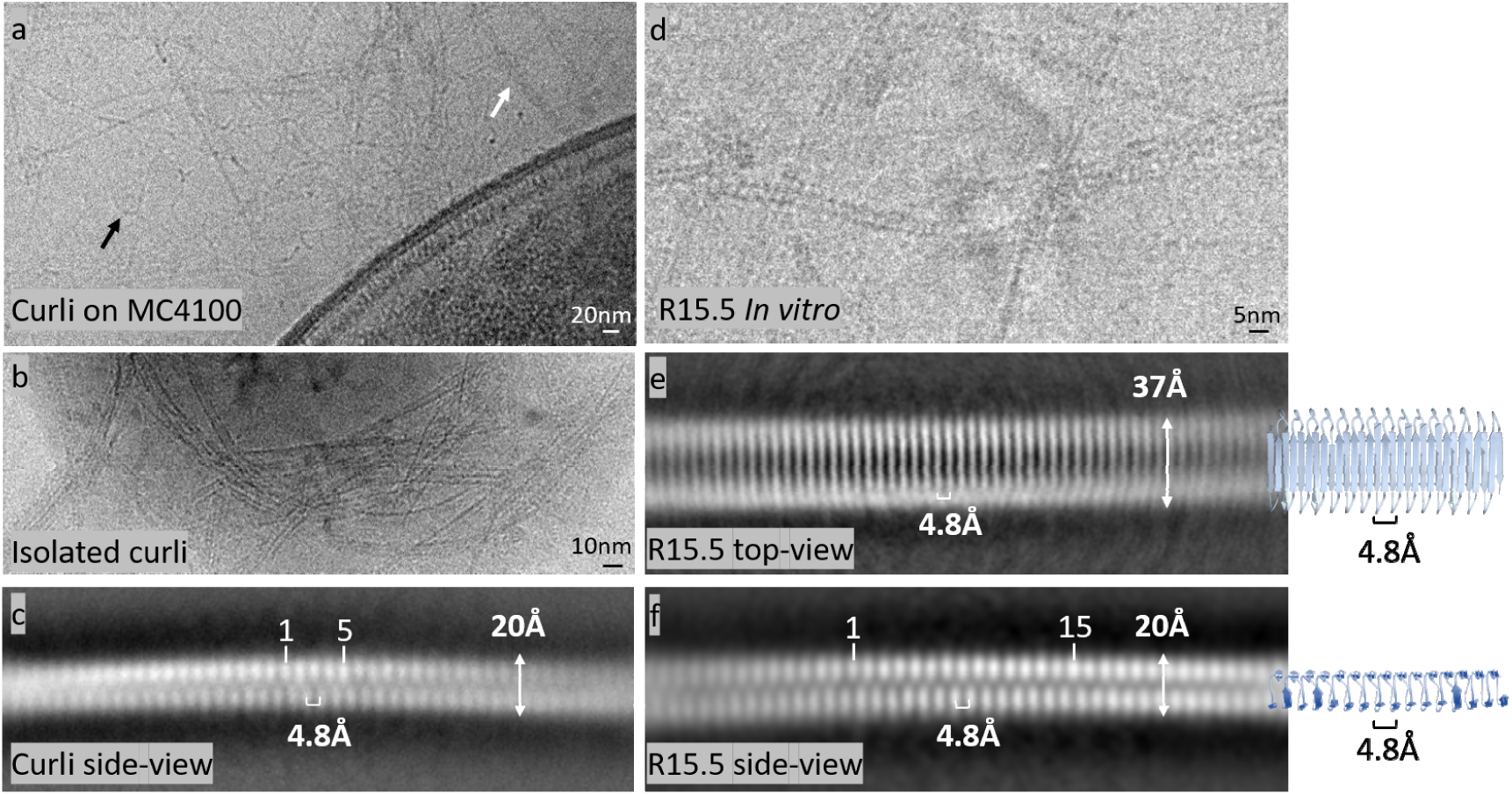
Cryo-EM reveals a conserved β-solenoid architecture. **(a)** low magnification (25k) cryoEM image of the extracellular matrix of MC4100 resolving two types of fibers: white arrow, curli filaments; black arrow, unidentified fibers, potentially eDNA or polysaccharide; **(b)** CryoEM image at 60k of isolated curli fibers; **(c)** side-view 2D class average of curli protofibrils; **(d)** CryoEM image at 60k of recombinant R15.5 fibers formed *in vitro* in 15mM MES 6.0; **(e)** and **(f)** top- and side-view 2D class averages of R15.5, respectively.

For R15.5, the head-to-tail mechanism of dimerization seems similar at first instance, i.e. β-solenoid augmentation via docking of open-ended sheets and complementarity of the steric zipper (Fig.3c; Supporting Figure 7). However, R15.5 has an uneven number of strands (i.e. 31), resulting in a motif A overhang both at N and C-termini. This geometry does not allow for simple docking of two molecules in the same orientation (Fig. 3a). Rather, a 180° rotation of the second molecule with respect to the first is required for a proper match of the single strand overhangs across the dimer interface – as is predicted by AF2 shown in Fig.3c and Supporting Fig.7. In this configuration, we again recover a near seamless transition between two molecules which is facilitated by the centro-symmetric nature of the steric zipper in this CS-class monomer. For this screw-axis R15.5 interface, 32 putative H-bonds are found which is remarkably consistent with the average of 29.6 between repeats within an R15.5 monomer. We also note that the second β-arc is reduced from 4aa to 3aa in the last two repeats. This in turn perfectly accommodates the continuation of the β-arcade 2 of the first monomer into β-arcade 1 of the second monomer across the interface.

Finally, we investigated heteromeric contacts between CsgA homologues from two different species. This is relevant because promiscuous cross-seeding has been shown to occur between curli produced in interspecies biofilms^35^. For this we looked at an AF2 model of a CsgA-CsgA *Citrobacter-Salmonella* dimer (pLDDT:79.8; pTMscore:0.78; Supporting Figure 8). This dimer is conceptually identical to the homotrimer shown in Fig.3a in that the R5 repeat of CsgA_Citrobacter docks onto the R1 repeat of CsgA_Salmonella, with no screw axis present. These results show that the conserved repeat architecture facilitates docking of disparate monomers into fibrils, thereby facilitating inter-species curlin cross-reactivity.

### CryoEM resolves the staggered, β-solenoid architecture of *ex vivo* and *in vitro* curli fibers

In their natural context, curli fibers are produced as an erratic, entangled mass that constitutes the major component of the extracellular matrix (ECM) under biofilm forming conditions ^18^. This is exemplified in the low magnification (1.88Å/pix; 20k; Fig.4a) cryoEM image we collected of the ECM produced by *E. coli* MC4100 after 72h of growth on YESCA agar at RT. In Fig4a, multiple filamentous structures can be discerned emanating from and engulfing a bacterial cell. Attempts to generate stable cryoEM class averages from digitally extracted filament segments were not successful at this stage. We therefore proceeded to isolate *ex vivo* curli fibers following the extraction protocol that was optimized by Chapman *et al*. ^36^. Extracted curli fractions were subjected to mild sonication (30sec; 10/10sec on/off pulses) to fracture and disentangle micron-sized curli conglomerates leading to a dispersed curli sample from which a cryoEM dataset was collected at 60k magnification (0.784Å/pix; Fig.4b). Although most curli still existed as large multi-filamentous bundles, single fiber fragments allowed 2D averaging using relion 3.1, yielding a unique “side-view” class average with secondary structure features present (Fig.4c). This revealed a cross-beta architecture characterized with 4.8Å repetition, a fibril width of 20Å, and a half unit stagger between opposing β-strands in the two sheets of theβ-solenoid. The corresponding power spectrum exhibits two broad maxima at 1/4.8Å^-1^ and 1/10 Å in a ∼13° and 90° angle to the meridian, reflecting, respectively, the staggered 4.8Å spacing of β-strands and the ∼10Å spacing of the two β-sheets (Supporting Fig.9). These numbers closely match the monomer and fiber architectures of EcCsgA predicted by AF2. The absence of other maxima in the power spectrum, suggest an absence of helical symmetry, although a low helical symmetry under the form of a two-fold screw axis cannot be ruled based on this data alone. However, our analysis of the AF2 trimeric structure suggests that the presence of a screw axis is unlikely, and also nanogold labelling patterns obtained by Chen et al. are in agreement with a purely translational head-to-tail propagation of CsgA subunits in curli fibers ^17^.

We also note that there are no clear features that delineate the interface between two successive CsgA monomers (cfr. a single monomer consists of 5 beta-arc-beta motifs), nor are there any low-resolution maxima in the power spectrum from which a fiber period could be estimated (Supporting Figure 9). This means that the lateral alignment of the curli segments in the classification protocol did not find a fibril register – a likely consequence of the quasi-seamless transition between successive monomers and the near-isomorphous nature of the repeats. Consequently, due to the lack of additional high resolution class averages corresponding to different fibril orientations, 3D reconstruction was not feasible at this point. Attempts to obtain further 2D classes using *in vitro* grown *Ec*CsgA fibers ^34^ were unsuccessful due to the high bundling tendency of the fibers.

In light of the colloidal instability issues of *Ec*CsgA, we decided to pursue the structure of a CsgA homologue. For this we selected R15.5 due to its high Asp and Glu content (20%; pI=2.98) leading us to hypothesize that R15.5 protofibrils would be less prone to assemble into larger structures because of stabilizing, repulsive electrostatics. R15.5 was recombinantly expressed and purified from inclusion bodies and left to polymerize after buffer exchange. TEM analysis revealed curli-like fibers that exhibited only minimal inter-fibril aggregation. Given its high Asp and Glu content, present in steric ladders on the R15.5 motif A (sheet 1) surface, we tested if R15.5 fibril formation could be tuned by changing pH or using Ca^2+^ as a counter ion to avoid charge repulsion. Somewhat unexpectedly, R15.5 still formed fibrils in the presence of 10mM EDTA, 15mM MES 6.0 (Supporting Fig.10) or in 50mM bicine pH 9.0 (Supporting Fig.10), conditions where formation of the Asp or Glu ladders upon R15.5 folding and polymerization were expected to result in charge repulsion. However, when we supplement 15mM MES 6.0 with 10mM CaCl_2_ (Supporting Fig.10) or change the buffer to 50mM Na-Acetate 4.0 (Supporting Fig.10) we observed a marked increase in fibrillar aggregates. These results demonstrate that the R15.5 amyloidogenicity is robust and to some extent tunable via solvent electrostatics.

In Fig.4d we show a representative cryoEM image of *in vitro* grown R15.5 fibrils and the corresponding 2D class averages, which we assign as “front” and “side” views. The R15.5 side-view is nearly identical to the side-view of *E. coli* curli, suggesting that *ex vivo* curli and *in vitro* R15.5 fibrils are structurally equivalent at the fold level and essentially in agreement with AF2 predictions. As was the case for EcCsgA, we detect no measurable twist along the fibril axis (Fig. 4f). Volume reconstruction was not feasible again, due to the pronounced preferential orientation in this dataset. Although initial tests showed that addition of CaCl_2_ can induce R15.5 aggregation -likely due to the formation of cross-fiber salt bridges-we prepared additional grids starting from a stock solution of R15.5 fibrils to which 1mM CaCl_2_ was added only seconds before deposition on a GO-coated grid and plunging in liquid ethane. The resulting dataset yielded class averages corresponding to additional, partially tilted, R15.5 orientations from which a low resolution C1 volume reconstruction could be attempted (Supporting Fig.11). The resulting 3D volume further supports the predicted β-solenoid fold but from which no further structural details can be determined.

### Higher order organization of curli fibers

We have found through structure prediction and experimental observation that the curli protofibril is formed of a highly regular supermolecular β-solenoid with an absent or negligible helical twist. Interestingly, apart from producing single protofilaments, curli fibrils had a tendency to form irregular or even systematic higher order structures (Fig. 5). EcCsgA curli, for example, frequently show a lateral association into thicker bundles, with average diameter of 17±9nm (mean, standard deviation, n=108) and outliers up to 48nm (Fig. 5g). R15.5 showed a range of lateral association, ranging from single fibrils (see above) over regular fibril dimers, to planar arrays wherein multiple fibrils are stacked side-by-side in an organized manner (Fig. 5a). 2D class averaging of boxed segments from such fiber dimers or fiber sheets resolves two or multiple parallel running R15.5 fibrils (Fig. 5b, c). For both fibrils in the fiber dimer or for at least the 3 fibrils in the center region of the fiber sheet, we clearly resolve the β-solenoid features. The strands of each fibril are aligned with the strands of the neighboring fibril which strongly suggests that they make specific inter-fibril contacts. If the inter-fibril packing was unspecific then one could expect the fibrils to ‘slide’ across each other which would yield a 2D class average wherein only a single fibril has its secondary structure features resolved. Such a situation is observed for EcCsgA fiber dimers, where one fibril is resolved at secondary structure level, whilst the partner fibrils show a smeared out projection in the 2D class averages (Fig. 5e). For R15.5 we hypothesize that the packing contacts between R15.5 fibrils are mediated via electrostatics. Specifically, the AF2 model of R15.5 predicts a square grid of glutamate and aspartate residues on the solvent exposed motif A flank, forming a large negatively charged patch (Fig.5d). As a result of the screw axis for an R15.5 fibril, that negative patch will alternate sides along the fibril axis, thereby facilitating the formation of salt-bridges between neighbouring fibrils mediated by divalent cations on either side of the fibril (Fig.5d). If we supplement the buffer solution with a stochiometric excess of Ca^2+^ ions -30mM CaCl_2_ added to 3µM R15.5-in the presence of 15mM MES pH 6.0 and 250µM potassium phosphate, then we no longer observe sheet formation, but rather encrustation of single R15.5 fibrils (Fig. 5e). TEM resolves a 2.5nm diameter R15.5 fibril at the core, enveloped by a thick crust that ranges from 10-30nm (Fig. 5f). We hypothesize that this crust is an inorganic deposit composed of a crystalline phase of calcium phosphate which nucleated on the D/E grid (Fig. 5c).

**Fig. 5:**
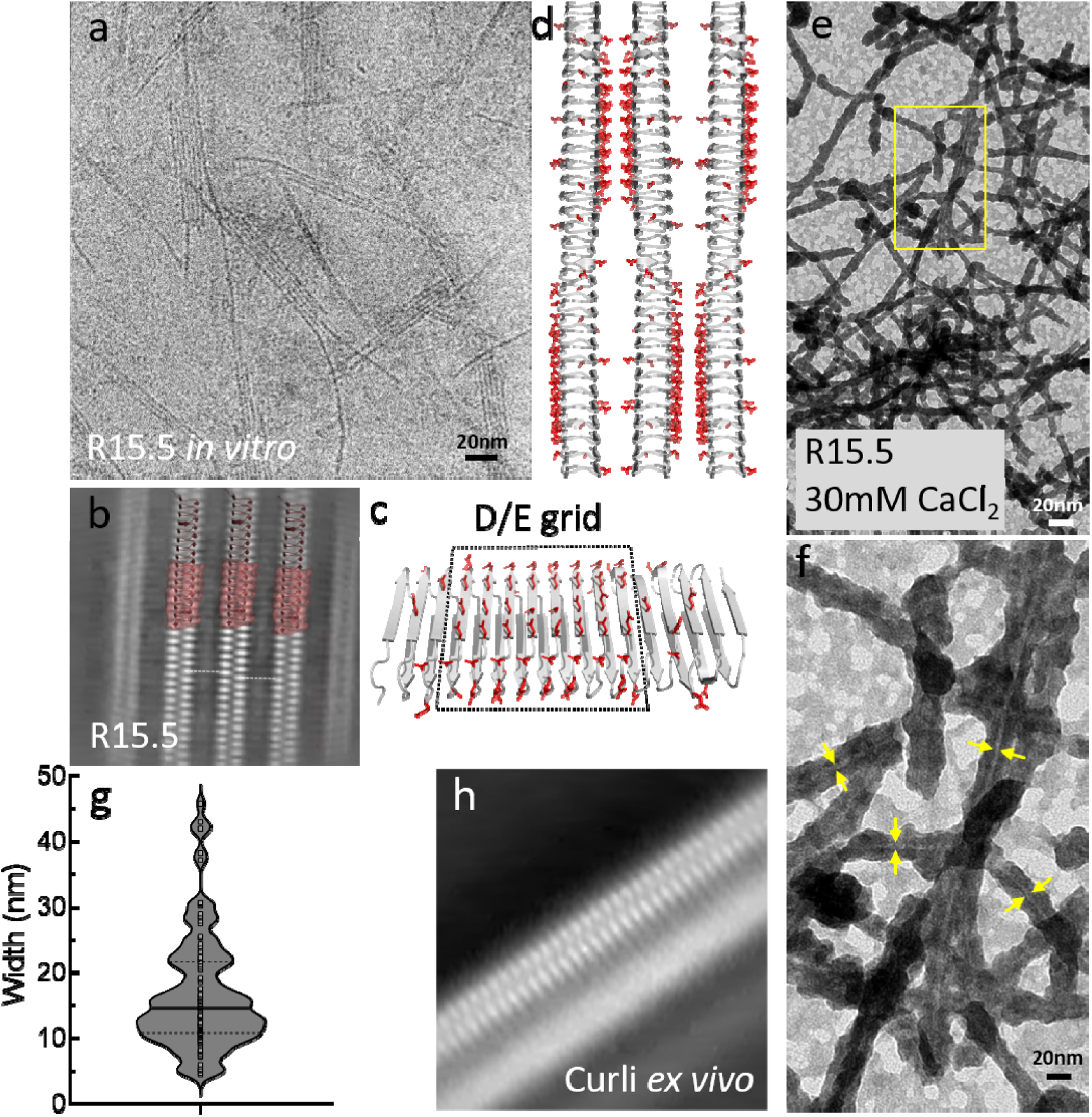
Hierarchical organization of curli fibers. **(a)** CryoEM image of side-way stacked, parallel R15.5 fibers organized into a planar array; **(b)** 2D class average of boxed segments of the core region of R15.5 arrays wherein the secondary structure of the three central fibrils is resolved; **(c)** Regular array of surface exposed asp and glu residues on the sheet 1 flank of the R15.5 monomer; **(d)** Idealized model for the supramolecular organization of R15.5 fibrils mediated by inter-fibril salt-bridges. The screw axis facilitates alternating binding interfaces between consecutive fibrils; **(e)** Encrustation of R15.5 fibrils in the presence of 30mM CaCl_2_ and trace amounts (±250µM) of KH_2_PO_4_/K_2_HPO_4_; **(f)** Zoom-in of the boxed area in (e) resolving the fibril core (yellow arrows) surrounded by a thick deposit; **(g)** Violin plot of the diameter of *ex vivo* curli fibers purified from the ECM of MC4100: solid line=median, dashed lines=quartiles; **(h)** 2D class average of curli fibers wherein only a single fibril with resolved secondary elements.

R15.5 could be conceptually similar in that regard to ice binding proteins ^37^ such as *Lp*IBP (3ULT) or *Tm*AFP (1EZG), in that it displays a regular array of acidic side chains on its surface, that may match with the lattice spacing of inorganic phosphate minerals, such as brushite or apatite. When present in the ECM of *Pontibacter korlensis* -which was isolated from the desert of Xinjiang in China ^38^-R15.5 could facilitate encrustation of the biofilm to help mitigate the adverse conditions (desiccation and UV-irradiation) that are associated with arid biotopes. This analysis is hypothetical at best, but it does raise the question regarding functional diversification and specialization of curli fibers across the Gram-negative microbiome.

## Discussion

Our structural understanding of amyloid fibers has advanced greatly in the last decade. A common denominator that has emerged from the extensive pool of experimental amyloid structures is the serpentine fold, i.e. a planar arrangement of beta strands and turns that form super-pleated ultra-structures that are stabilized by steric zipper interactions and strand-strand docking. Strikingly, amyloid fibers can exhibit a large degree of polymorphism at the level of the serpentine fold, and/or at the level of protofibril contacts ^2-4^. Small changes in the primary sequence or in the polymerization conditions can lead to drastic changes in the final ultrastructure of the amyloid fiber. That sensitivity of the quaternary structure to initial conditions can likely be attributed to the fact that for most of the reported peptides/proteins there has been no (or perhaps even a negative) evolutionary pressure to fold into a specific amyloid structure. Rather, most characterized amyloids are formed as an unintended consequence of an environmental trigger leading to an amyloidogenesis process that is stochastic and susceptible to external perturbations.

This stands in sharp contrast to functional amyloids that have been formed by evolutionary processes, and for which uncontrollable polymorphism may well be biologically intolerable. This indeed appears to be the case for curli. After more than two decades of research on curli, there are no experimental observations of fiber polymorphism, i.e. there has been consistent reporting of protofibril diameters and no measurable helical symmetry or changes thereof. Equally, we find no evidence for even sparsely populated fiber class averages corresponding to a helical curli protofibrils. For curli, structural consistency is likely a necessity that is imposed by the tethering mechanism to the extracellular CsgG-CsgF-CsgB complex. In a nucleation-dependent fibrillation process, structural conformers could be imposed by the CsgG-CsgF-CsgB complex. However, we find the protofibrils of *ex vivo* isolated and *in vitro* formed curli to comprise an identical β-solenoid structure, which strongly suggests that this structure is an intrinsic property of the CsgA sequence. Additionally, Zhou and coworkers have shown that promiscuous cross-seeding can occur between curli produced in interspecies biofilms – a process that is likely dependent on a conserved curli architecture ^35^. For curli, the structural consistency exists in the form of a conserved β-solenoid fold of the monomeric building blocks. Notwithstanding subtle variations in the β-solenoid scaffold and its local ‘decorations’ under the form of insertions, AF2 predictions are remarkably conservative despite large variations in primary sequence. Open ended β-solenoids that lack capping domains give rise to a polymerization mechanism that relies mostly on the compatibility of secondary structure motifs, i.e. β-sheet augmentation and β-arcade extension which is mediated predominantly by main-chain contacts, reliant only to a lesser extent on local sidechain contacts between chains. Additional structural preservation and complementarity is guaranteed by a high degree of conservation of a limited number of key residues that partake in steric zipper contacts and the stabilization of β-arcade structures. These structural insights provide a mechanistic understanding of inter-species curli cross-seeding, as well as a molecular model for curli tethering to the cell via CsgB. Extracellular matrix components such as curli have been proposed as a secreted public good, at least in single species biofilms ^39^. The structural conservation in curli subunits may enable mixed fiber formation and thus allow multispecies contribution of curli monomers to the biofilm matrix.

Curli fibers carve out a unique niche in the superfamily of amyloid proteins. Curli fibers do not show the serpentine fold typically observed for pathological amyloids or recently seen in peptide-based functional amyloids such as LARK-like amphibian antimicrobials ^15^. CsgA marries characteristics of globular proteins with features that are often associated with amyloids such as a stacked cross beta structure and nucleation-dependent polymerization. On the one hand, curli seeds form during an initial stochastic phase of nucleation, followed by a stage of step-wise and self-catalyzed extension of cross-β fibers that are extremely robust ^34^. On the other hand, however, MSA analysis uncovers strong co-evolutionary couplings that provide distance restraints that almost deterministically predict a unique, folded, monomeric state. In that regard, curli formation can also be viewed as a classical polymerization process with the added complexity that the rate of folding is likely commensurable to the rate of aggregation, and that folding is templated once primary nucleation has taken place. In this model, the single-strand overhang at the fiber extremities would provide a templating surface that recruits and helps fold incoming CsgA protomers. At least in vitro, EcCsgA curli fibers extend from both fiber ends, albeit with markedly different kinetics ^34^. Future studies will be needed to determine the structural underpinning of these polar growth kinetics, and determine the orientation of curli fibers on the cell surface.

Finally, we make liberal use of the steric zipper terminology to refer to the interlacing pattern of inwards facing, conserved residues, but for CsgA that inter-digitation does not occur in the same plane -as is classically the case for PAs-due to the stagger between the opposing strands. It is therefore clear that curli blur the lines between classical, protein polymers and amyloid fibers, prompting a re-evaluation of the definition of a functional amyloid.

## Author contributions

MS and HR designed the project. All authors contributed to cryogenic freezing, cryoEM imaging and data processing. MS wrote the manuscript with contributions from all authors.

## Data Availability Statement

The AF2 predictions discussed in main text figures 2 and 3 are supplied as supplementary data to this work. Additional data is available from the corresponding authors upon reasonable request.

## Acknowledgements

We thank Marcus Fislage and Dirk Reiter at the VIB-VUB Facility for Bio Electron Cryogenic Microscopy (BECM) and for assistance in data collection. We thank Jolyon Claridge for assistance in the genome mining. This work was funded by VIB, EOS Excellence in Research Program by FWO through grant G0G0818N to HR and G043021N to MS.

## Supporting Figure

**Supporting Figure 1:**
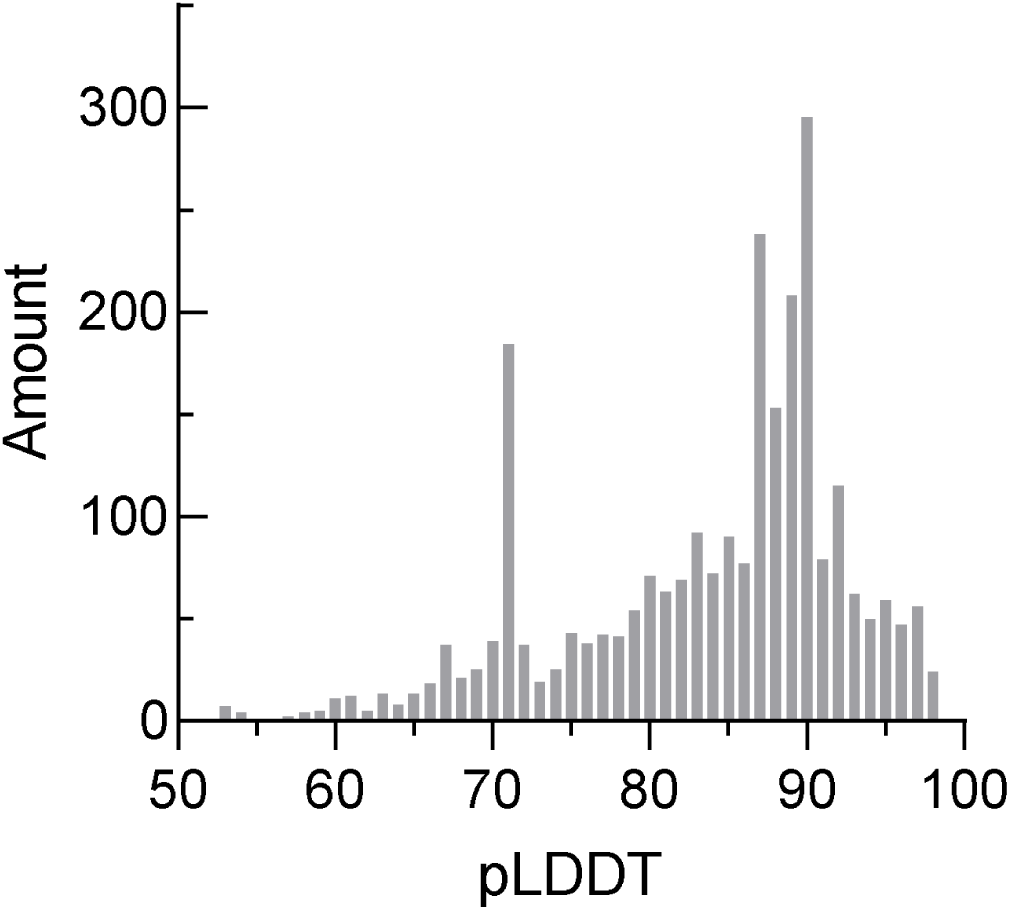
Histogram of the reported pLDDT values for the total dataset of predicted CsgA structures. DeepMind reports pLDDT > 90 as high accuracy predictions, between 70 and 90 as good backbone predictions, and pLDDT < 70 as low confidence and to be treated with caution.

**Supporting Figure 2:**
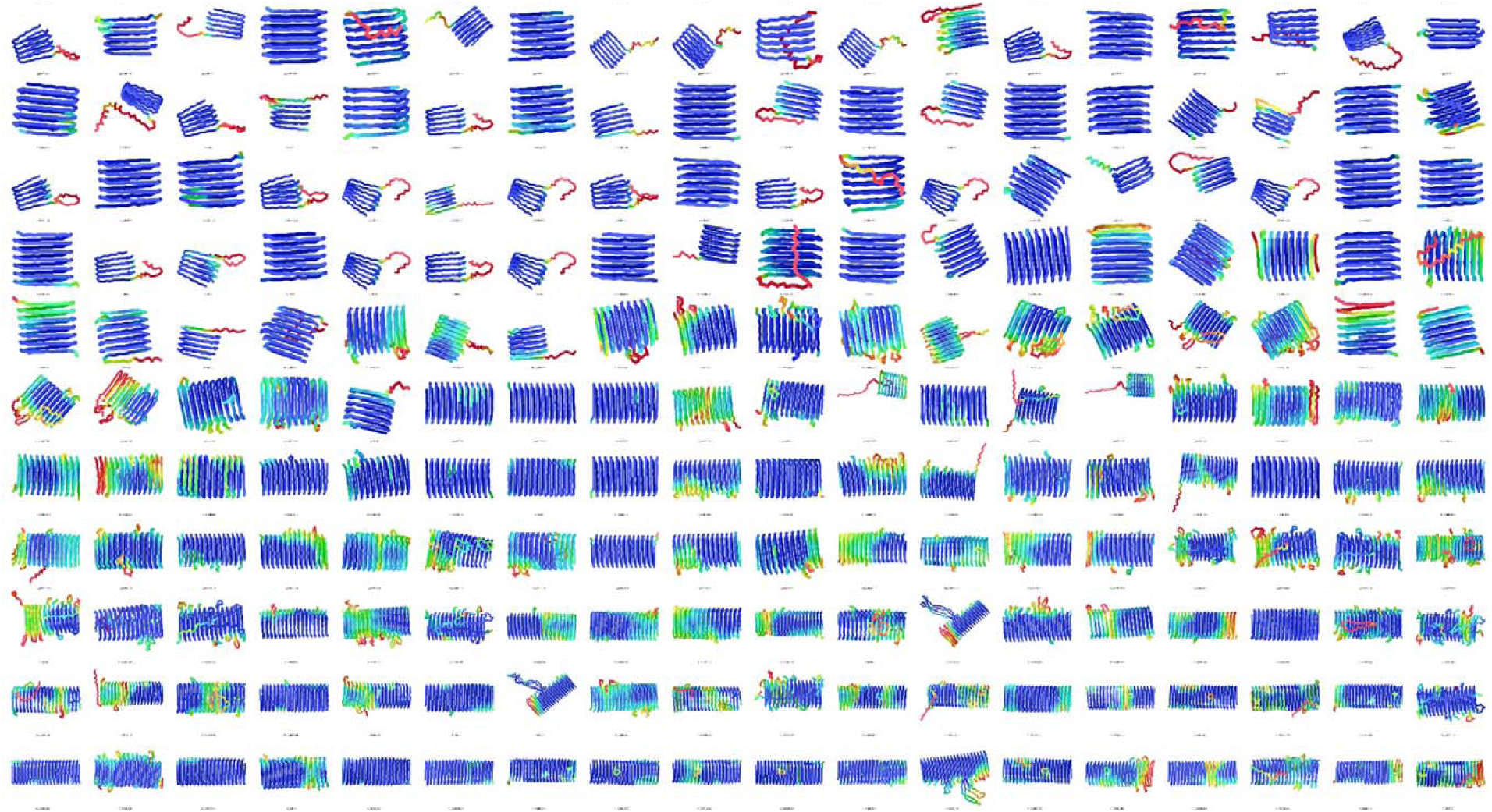
Representative collage of a subset of CsgA homologue models produced by AlphaFold2, covering the structural diversity in the database. Colour coding according to pLDDT values.

**Supporting Figure 3:**
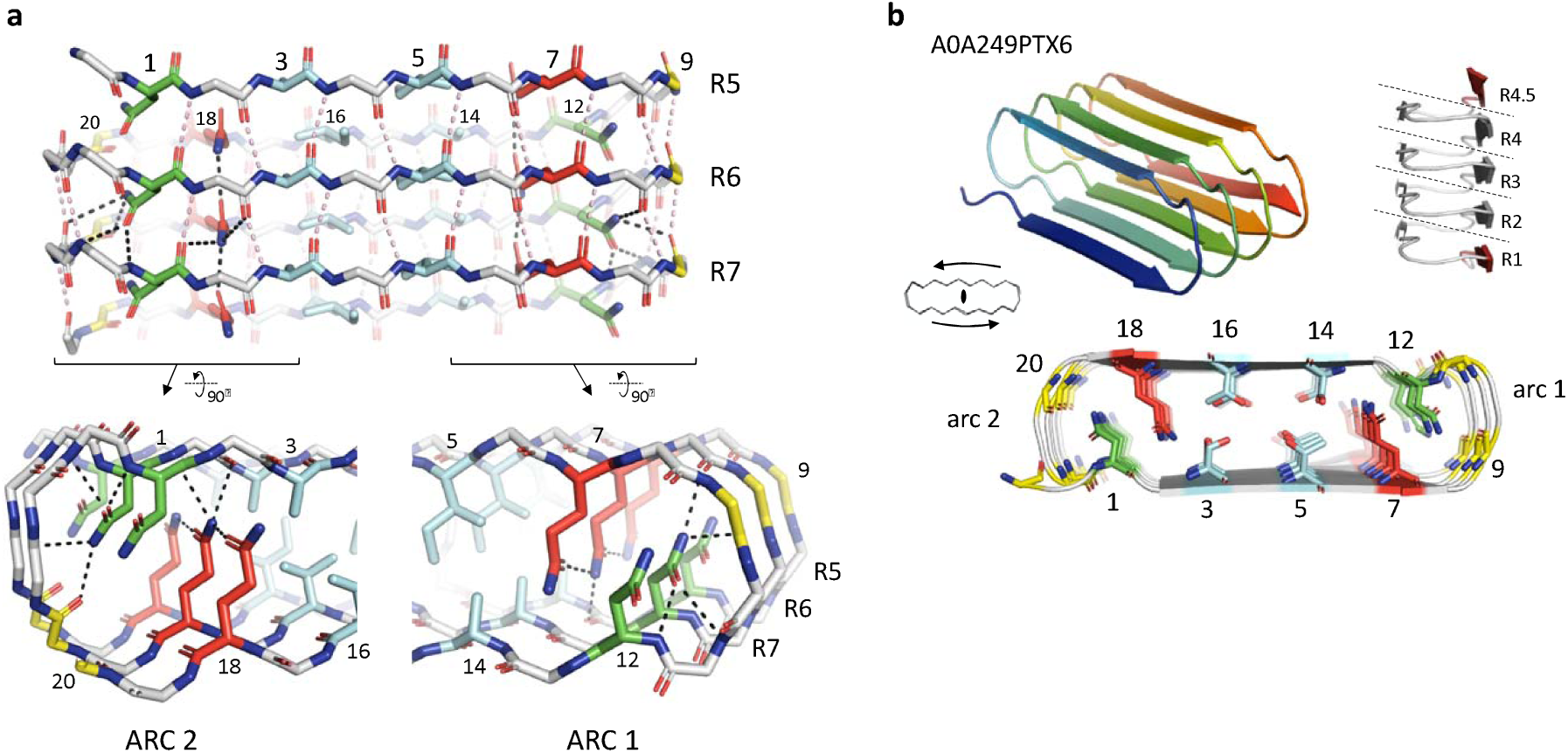
Structural analysis of curlin repeat contacts. **(a)** Stick representation of three representative curlin repeats (R5-R6-R7) of *Pontibacter korlensis* CsgA (i.e. R15.5) shown in frontal view (top, with motif A facing forward), or in a close-up axial view of β-arc 1 (lower right) and β-arc 2 (lower left). For clarity, sidechains of surface exposed residues are not shown. N, Q and Ψ in motif A and B are colored green, red and sky blue, resp. Gly are colored yellow. Main chain H-bonds between consecutive curlin repeats are shown as pink dash. H-bonds formed by N, and Q in the curlin repeats are colored black (shown for R6 only). **(b)** Predicted structure of CsgA-like protein A0A249PTX6 of *Sinorhizobium fredii* shown in ribbon representation (upper left, blue to red from N-to C-term), in axial view (bottom), with inward facing residues shown in stick representation and colored as in (a), or in side view (upper right), with dashed lines indicating consecutive curlin repeats (labeled R1 to R4, terminating in half-repeat R4.5).

**Supporting Figure 4:**
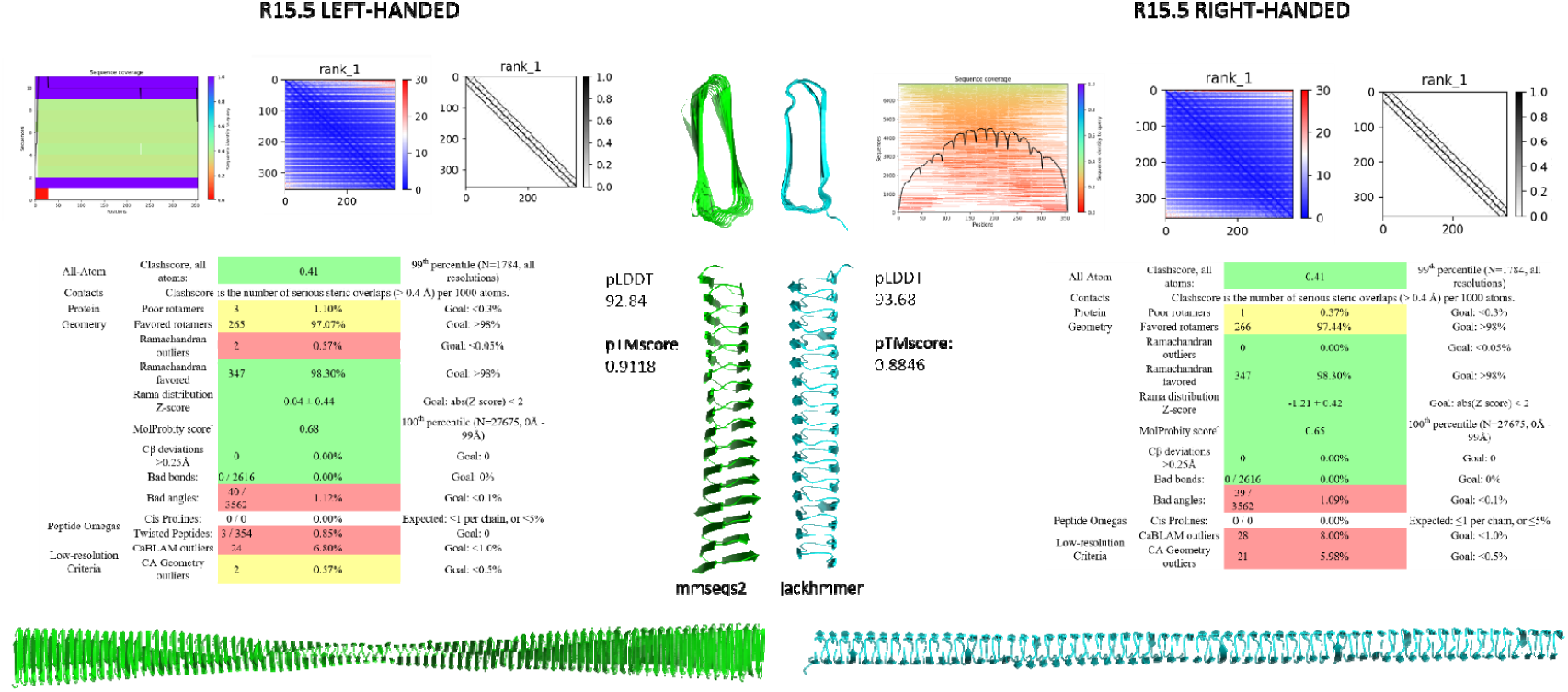
Structural analysis of AMBER relaxed, left- and right-handed AF2 models of R15.5. Panels from left to right as produced by ColabFold (https://colab.research.google.com/github/sokrypton/ColabFold/): MSA coverage; predicted alignment error; Predicted contacts. Tables: MolProbity report for each model taken from http://molprobity.biochem.duke.edu/. The AF2 pLDDT and pTMscores for the left- and right-handed model are nearly identical, as well as the contact map and alignment error matrix. Molprobity scores for both structures are excellent, i.e. 0.68 and 0.65 (100^th^ percentile of the reference database) and nearly identical. The only meaningful difference between both models -apart from the chirality of the fold-is the twist of the of the β-solenoid. The right-handed R15.5 has no measurable twist, whereas the left-handed model has a 17° twist from N-to C-terminus. Simulated fibril models based on left- or right-handed protomers are shown below.

**Supporting Figure 5:**
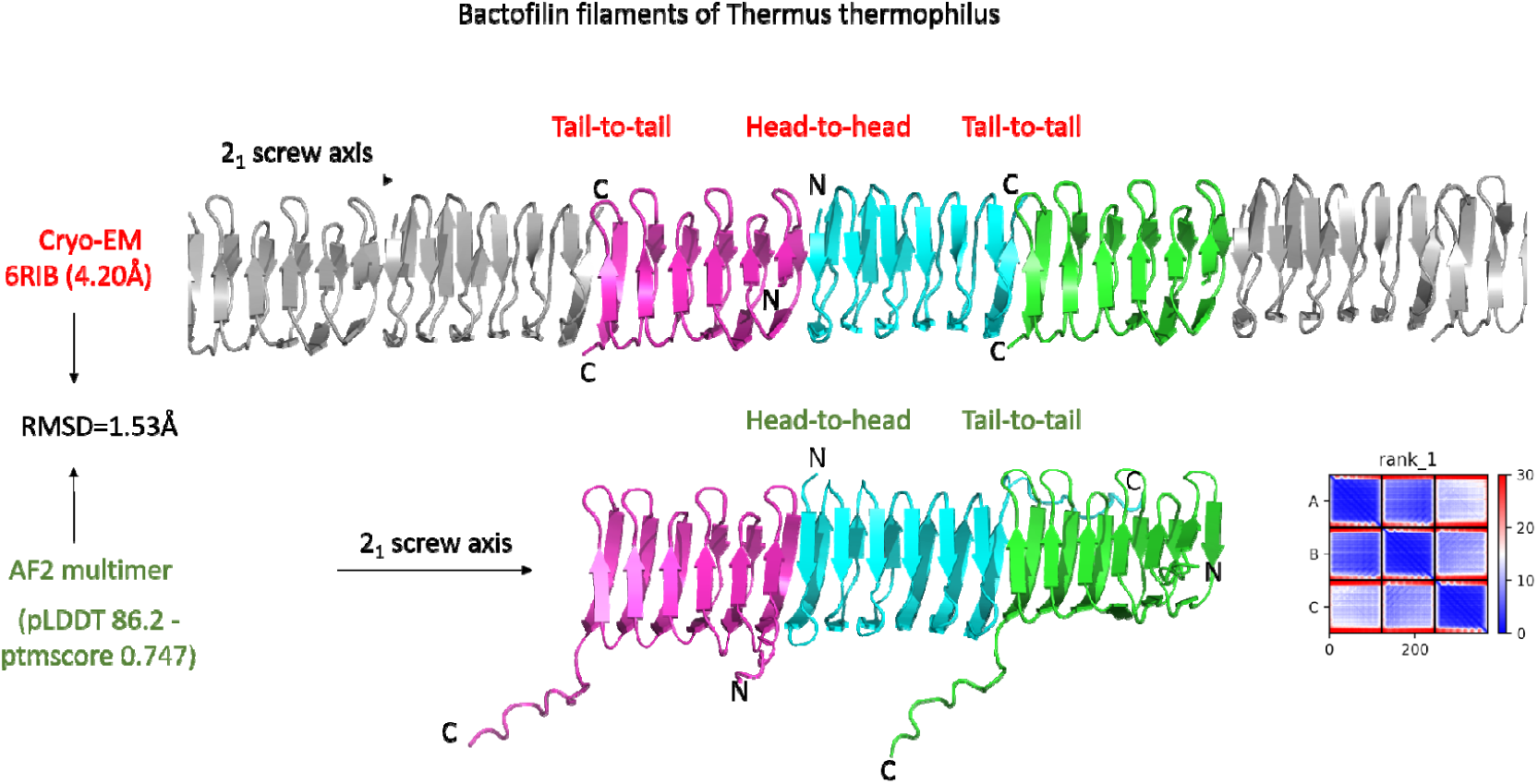
Benchmarking example of the proposed AF2 methodology to predict filament structures: we compare an experimental cryo-EM structure (Bactofilin filaments of *Thermus thermophilus*) with a trimer that was predicted using AF2 multimer. Overall, there is excellent agreement between the predicted and experimental structure, with an RMSD of 1.53Å. AF2 accurately predicts the apolar nature of the filaments (i.e. succession of head-to-head and tail-to-tail interfaces) as well as the helical nature (i.e. screw axis). Note that the cryoEM structure was deposited on 2019-04-23, whereas AF2 is trained on protein chains in the PDB released before 2018-04-30.

**Supporting Figure 6:**
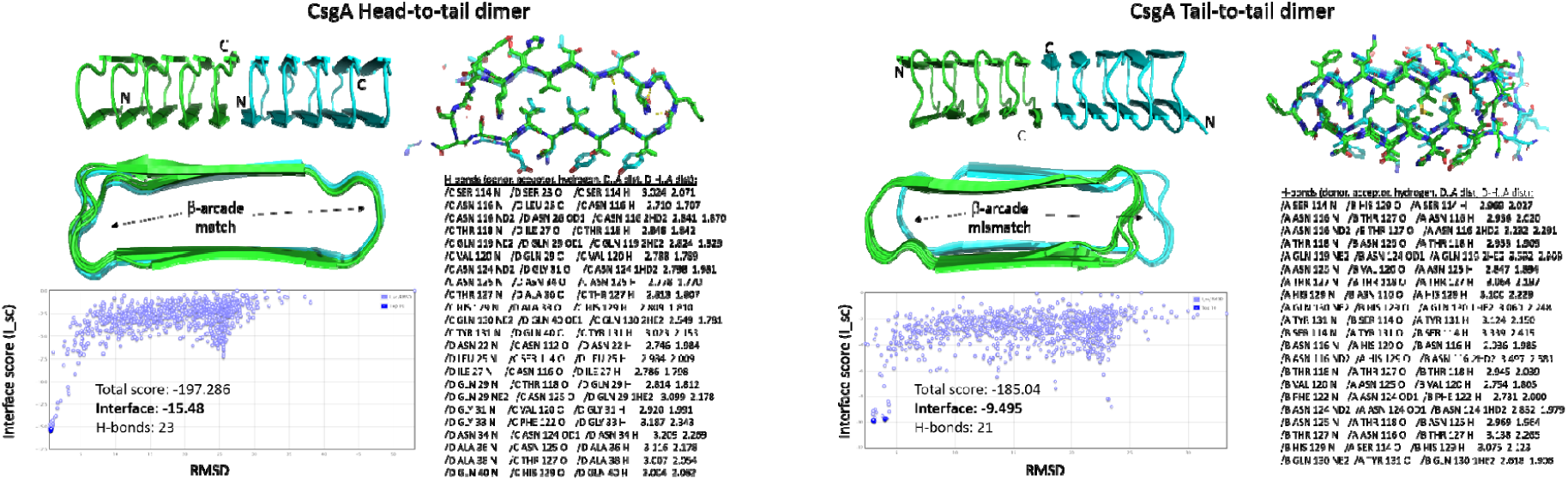
*In silico* models of a head-to-tail and tail-to-tail CsgA dimer. The tail-to-tail dimer model was generated by initial manual placement of two CsgA AF2 monomers (without N_22_) in a ‘tail-to-tail’ configuration, followed by local docking using RosettaDock. For the head-to-tail dimer we used the AF2 dimer model as an input structure for RosettaDock. Both models shown here correspond to their respective top-ranking structures that were generated by the ROSIE server. We plot the interface score (I_sc) for 1000 randomly placed models and show the total score, I_sc value and number of putative H-bonds for the top ranking models, respectively. The head-to-tail dimer packs in a parallel fashion resulting in β-sheet augmentation, and uninterrupted matching of the β-arcades, whereas the tail-to-tail dimer packs anti-parallel, triggering a lateral offset of 1 residue between both strands at the interface, and poor contacts at the arcades as a result. Putative H-bonds were detected using ChimeraX v1.2 using relaxed settings.

**Supporting Figure 7:**
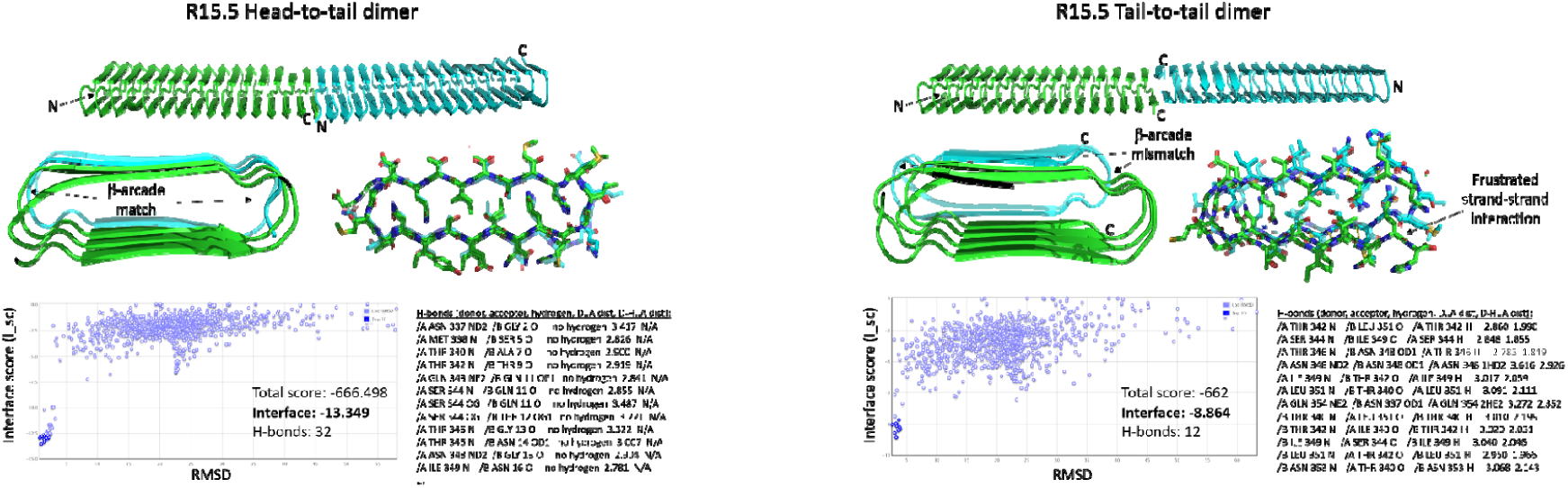
*In silico* models of a head-to-tail and tail-to-tail R15.5 dimer. The tail-to-tail dimer model was generated by initial manual placement of two R15.5 AF2 monomers in a ‘tail-to-tail’ configuration, followed by local docking using RosettaDock. For the head-to-tail dimer we used the AF2 dimer model as an input structure for RosettaDock. Both models shown here correspond to their respective top-ranking structures that were generated by the ROSIE server. We plot the interface score (I_sc) for 1000 randomly placed models and show the total score, I_sc value and number of putative H-bonds for the top ranking models, respectively. The head-to-tail dimer packs in a parallel fashion resulting in β-sheet augmentation, and uninterrupted matching of the β-arcades. Similarly to the tail-to-tail dimer for CsgA, the tail-to-tail R15.5 dimer packs anti-parallel, triggering a lateral offset of 1 residue between both strands at the interface, and poor contacts at the arcades as a result. Putative H-bonds were detected using ChimeraX v1.2 using relaxed settings.

**Supporting Figure 8:**
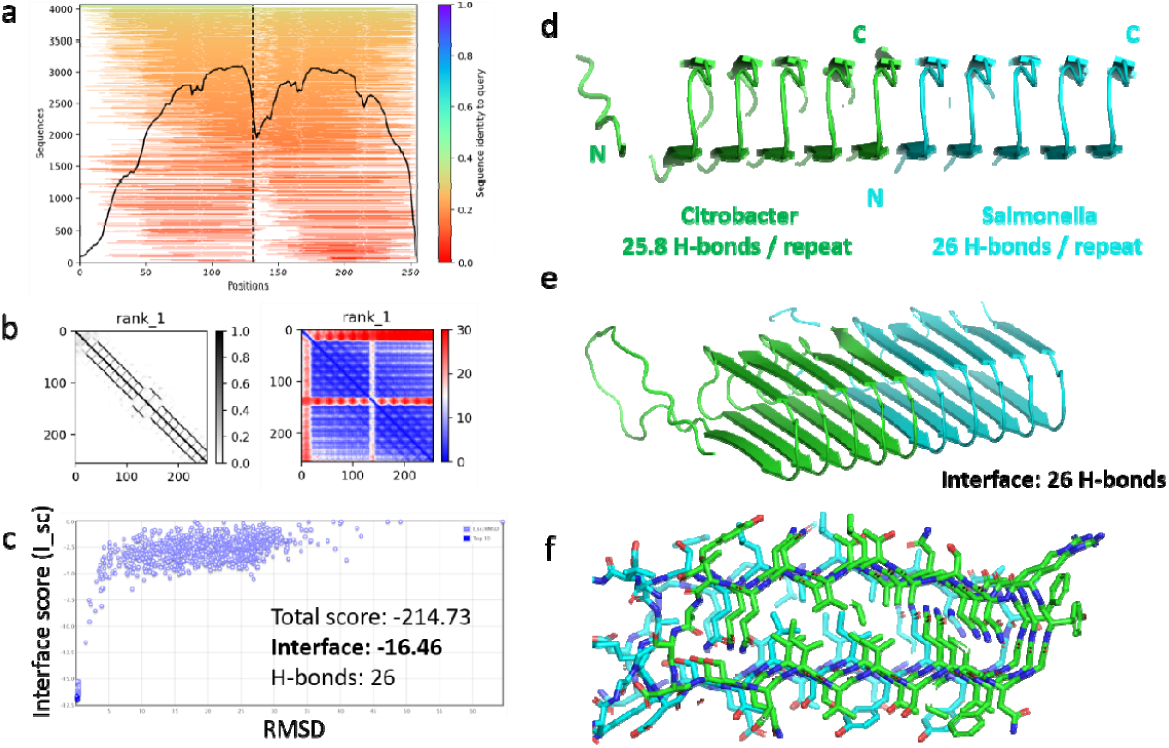
Predicted heteromeric dimer of CsgA monomers from *Citrobacter* and *Salmonella*. **(a)** Multiple sequence alignment; **(b)** Predicted contacts and alignment error; **(c)** Rosettadock interface score as a function of the RMSD with respect to the AF2 input structure; **(d)** Side-view and **(e)** tilt-view of a head-to-tail heteromeric dimer of CsgA-CsgA *Citrobacter*-*Salmonella* as predicted by AF2. Each respective CsgA monomer has 25.8 and 26 predicted H-bonds per repeat on average. The interface between both subunits entails 26 putative H-bonds; **(f)** On-axis view in stick representation highlighting the uninterrupted steric zipper columns across the dimer interface.

**Supporting Figure 9:**
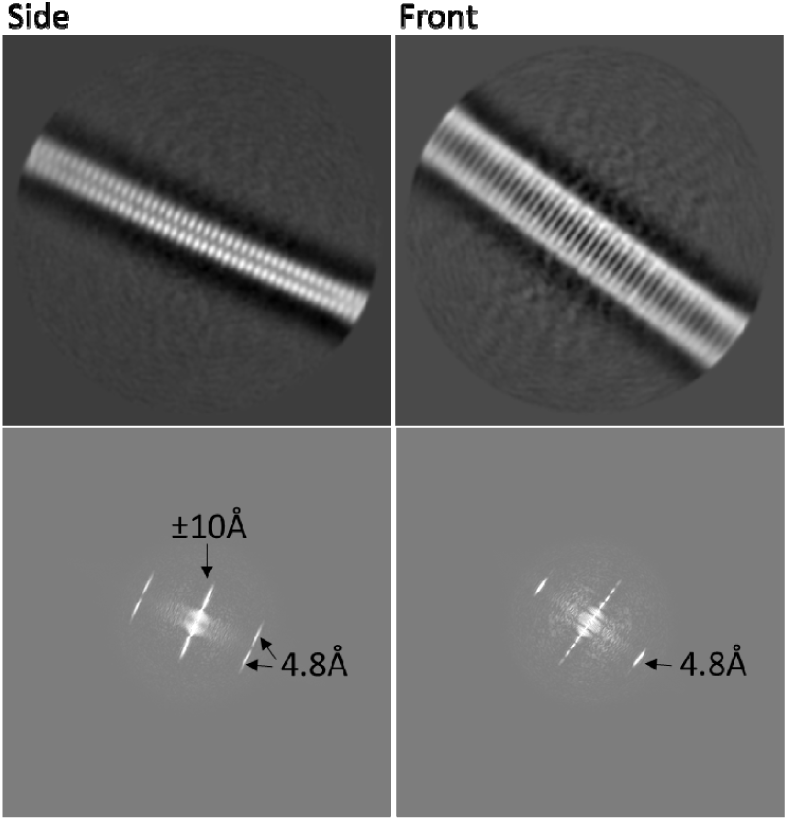
Side and front view 2D class averages (box-size: 300 × 300 pixels at 0.784 Å/pix) and corresponding power spectra output by Relion 3.1.3 at a sigma contrast value of 5.0.

**Supporting Fig 10:**
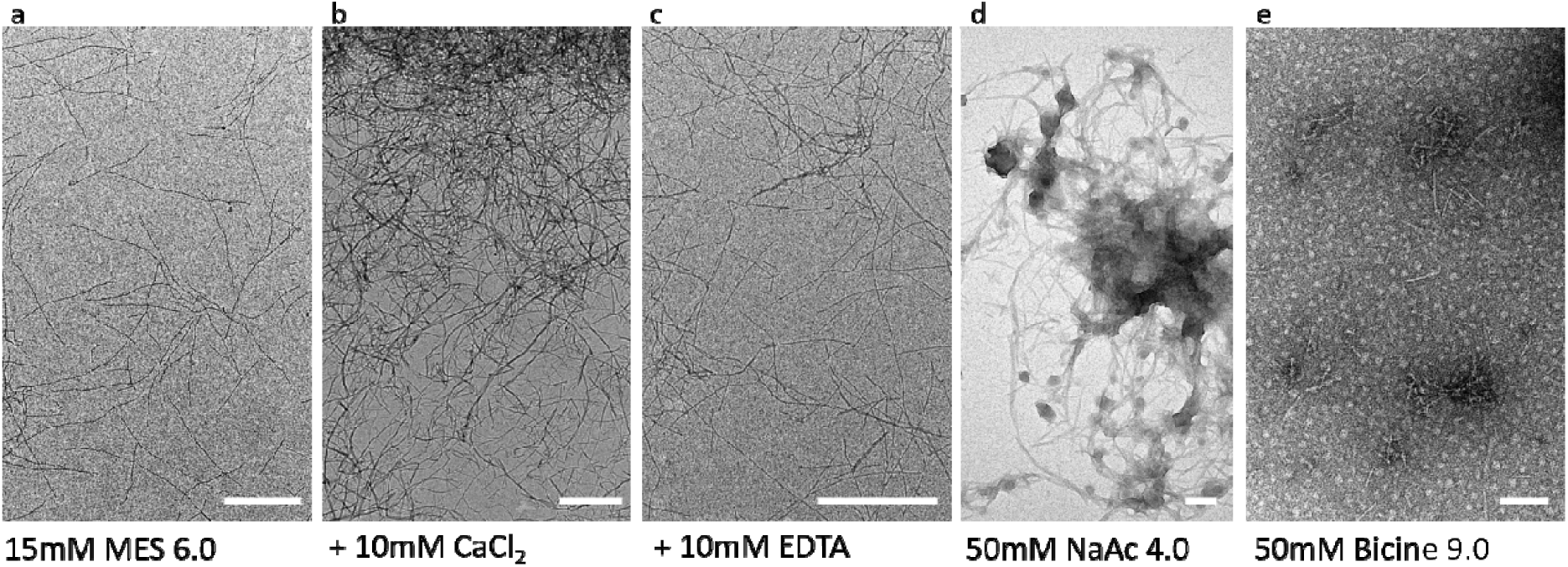
Negative stain TEM images of R15.5 fibrils in **(a)** 15mM MES 6.0; **(b)** 15mM MES pH 6.0 with 10mM CaCl_2_; **(c)** 15mM MES 6.0 with 10mM EDTA; **(d)** 50mM NaAc pH 4.0; **(e)** 50mM Bicine pH 9.0. R15.5 monomer (5µM) stocks in 8M urea were desalted using a zeba spin column to the corresponding buffer and incubated at RT ON.

**Supporting Figure 11:**
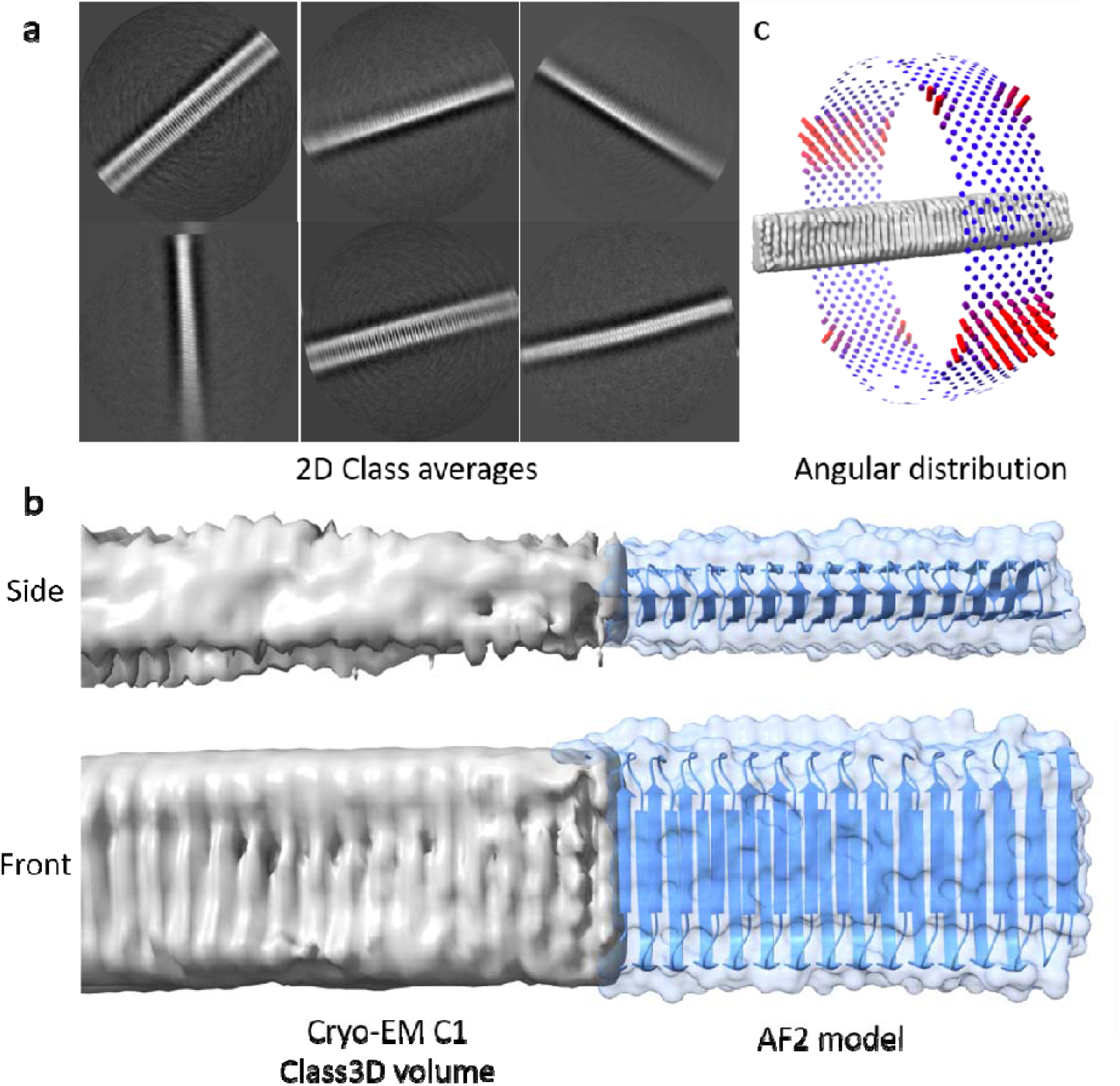
**(a)** 2D class averages of R15.5 fibrils in the presence of 1mM CaCl_2_. For this, R15.5 fibrils (in 15mM MES 6.0) were supplemented with 1mM CaCl_2_ (final concentration) seconds before deposition on a GO-coated grid and plunging in liquid ethane; **(b)** Corresponding reconstructed volume (Relion 3.1; Class3D; C1) contoured at level 0.00129; **(c)** Heatmap of the orientational distribution revealing preferential orientation.

## Materials and Methods

### Bacterial genome mining and AlphaFold2 modelling

To search for CsgA/B homologs, we set up a local Refseq genome database (ftp://ftp.ncbi.nlm.nih.gov/genomes/refseq/bacteria/). Genome searching was done using HMMER v3.3.2 with a threshold value of 1e-5, using the curated profile hidden Markov models from Dueholm *et al* ^19^. N-terminal leader sequences were removed using SignalP_6.0, and from this set of mature sequences duplicate entries were deleted. Curlin repeat sequences were extracted using a series of consecutive regular expression searches. First, we searched for **X**_**6**_**QX**_**10**_**Q** motifs (regex: .{6}Q.{10}Q), then we iteratively searched for **NX**_**5**_**QX**_**9**_ motifs (minimal 22 residues) that lack the second Q, but have an N at position 1 (regex: ([a-z]{22,})(N.{5}Q.G.{9,})). In a third step we also included longer, degenerate motifs **X(AIVLSTG)X(IVLQTA)xQxGx9**, for which we used the following regular expression: ([a-z]{22,})(..[AIVLSTG].[IVLQTA].Q.G.{9,}). All the extracted curlin repeat sequences were compiled into a single multi-fasta, and sequence logos were created using Weblogo 3 (http://weblogo.threeplusone.com/create.cgi).

Batch protein structure prediction of the homolog dataset was done using the localcolabfold implementation ^30-32^ (https://github.com/YoshitakaMo/localcolabfold) of AlphaFold2 using a tolerance value of 0.5, running for 6 recycles and outputting 5 models. Models were ranked using the pLDDT score. Models corresponding to figures 2a, 2b, 3 were produced using the AlphaFold2_advanced.ipynb Jupyter notebook at https://github.com/sokrypton/ColabFold using Jackhmmer, a tolerance value of 0.1, 24 recycles and outputting 5 models, and retaining the top-ranked models. For multimeric predictions, primary sequences of different subunits were concatenated and separated by a ‘/’.

### Protein production and purification

CsgA (P28307) and R15.5.*5* (A0A0E3UX01) were cloned into pET22b via the NdeI site without their signal sequence but with a C-terminal tag 6xHis-tag. Expression was induced in BL21(DE3) ΔslyD cells by addition of 1 mM IPTG after an OD600nm of 0.6 was reached. Cells were harvested by centrifugation at 5,000 g for 10 minutes after one hour of induction. Pellets were lysed for 30min in buffer A (50 mM Kpi pH 7.2, 500mM NaCl, 8 M urea, 12.5mM imidazole) and the cell lysate was centrifuged at 40,000 g for 30 minutes at 20°C. After sonication to reduce the viscosity of the lysate, the supernatant was loaded on a HisTrapTM FF column (GE Heathcare Life Sciences) equilibrated in 5 column volumes (CV) of buffer A. After washing in 10 CV buffer A, the protein was eluted using buffer B (50 mM Kpi pH 7.2, 8 M Gnd HCl, 250 mM imidazole). Relevant protein fractions were pooled and filtered with a 0.22µm cutoff filter to remove any potential amyloid seeds and stored at – 80°C.

To prevent any unwanted amyloid formation in our protein stock solutions, all purification steps were performed under denaturing conditions (8 M urea) and handling times at room temperature were reduced to an absolute minimum. This approach allows us to store CsgA and R15.5 in their pre-amyloid, unfolded form and gives control over the exact starting point of polymerization by buffer switching to native conditions, i.e. 15 mM MES pH 6.0. To remove urea, ZebaTM Spin Desalting columns (7K MWCO) (Thermo Scientific) or 5mL HiTrap Desalting Columns (GE Healthcare) were used.

### Negative stain TEM grid preparation

Negative stain TEM (nsTEM) imaging of R15.5 filaments was done using formvar/carbon-coated copper grids (Electron Microscopy Sciences) with a 400-hole mesh. The grids were glow-discharged (ELMO; Agar Scientific) with 4 mA plasma current for 45s. 3 µl of *in vitro* assembled R15.5 filament solution was applied onto the glow-discharged grids and left to adsorb for 1 minute. The solution was dry blotted, followed by three washes with 15 µl Milli-Q. After that, grids were dipped into 15 µl drops of 2% uranyl acetate three times for 10 seconds, 2 seconds, and 1 minute respectively, with a blotting step in between each dip. The excess stain was then dry blotted with Whatman type 1 paper. All grids were screened with a 120 kV JEOL 1400 microscope equipped with LaB6 filament and TVIPS F416 CCD camera.

### Cryo-EM grid preparation and image acquisition

High-resolution cryo-EM datasets were collected using in-house Graphene Oxide (GO) coated Quantifoil™ R2/1 400 copper mesh holey carbon grids. For GO coating, grids were glow-discharged at 5 mA plasma current for 1 minute in an ELMO (Agar Scientific) glow discharger. 3 µL of 0.6 mg /mL (GO) solution was then pipetted onto the grids and incubated at room temperature for 2 minutes. The GO solution from the grids was blotted out with a Whatman grade 1 paper after which it was washed with 20 µL of filtered Milli-Q water. The GO-coated grids were left to dry for 20 minutes. Gatan CP3 cryo-plunger set at room temperature and relative humidity of 90 percent was used to prepare the cryo-samples. 3 µL of filament solution was applied on the GO grid and incubated for 30 seconds. The solution was dry-blotted from both sides using Whatman type 2 paper for 3 seconds with a blot force of 0 and plunge-frozen into precooled liquid ethane at -176 °C. High resolution cryo-EM 2D micrograph movies were recorded at 300 kV on a JEOL Cryoarm300 microscope equipped with an in-column Ω energy filter (operated at slit width of 20 eV) automated with SerialEM 3.0.8 ^40^. The movies were captured with a K3 direct electron detector run in counting mode at a magnification of 60k with a calibrated pixel size of 0.764 Å/pix, and exposure of 64.66 e/Å^2^ taken over 61 frames. In total 3960 and 4455 movies were collected within a defocus range of 0.5 to 3.5 micrometers for *ex-vivo* curli and R15.5, respectively.

### Image processing

All the dose-fractioned movies were corrected for beam-induced motion using MOTIONCORR2 ^41^ implemented in RELION 3.1^42^. The contrast transfer function (CTF) of the motion-corrected images were calculated using CTFFIND4 ^43^. The R15.5 filaments were manually boxed using *e2helixboxer*.*py* of the EMAN2 package ^44^ while the coordinates of the *ex vivo* curli were auto-picked by SPHIRE-crYOLO using a model trained on 1000 manually picked filaments. The filaments were segmented and extracted into particles of box size of 300 pixels in RELION 3.1 which resulted in 341691 and 414352 particles of *ex vivo* curli and R15.5 respectively. To filter out non-ideal particles, multiple rounds of 2D classification were run in RELION 3.1 with a regularization parameter T value of 10 with each run consisting of 50 iterations. Several rounds of filtering resulted in datasets of 41,806 and 84729 enriched particles of ex *vivo* curli and R15.5 respectively. None of the resulting 2D class averages showed any signs of helical nature as judged from the peaks of the Fourier transform of the filament images. Our attempts to find helical parameters by optimizing the regularization parameter T and box size did not lead to improved 2D classes. 3D volumes R15.5 were generated in RELION 3.1 taking an empty cylinder volume with a diameter of 4nm as initial model.

